# Thermodynamic impacts of combinatorial mutagenesis on protein conformational stability: precise, high-throughput measurement by Thermofluor

**DOI:** 10.1101/591495

**Authors:** Violetta Weinreb, Gabriel Weinreb, Srinivas Niranj Chandrasekaran, Jhuma Das, Nikolay V. Dokholyan, Charles W. Carter

## Abstract

The D1 switch is a packing motif, broadly distributed in the proteome, that couples tryptophanyl-tRNA synthetase (TrpRS) domain movement to catalysis and specificity, thereby creating an escapement mechanism essential to free-energy transduction. The escapement mechanism arose from analysis of an extensive set of combinatorial mutations to this motif, which allowed us to relate mutant-induced changes quantitatively to both kinetic and computational parameters during catalysis. To further characterize the origins of this escapement mechanism in differential TrpRS conformational stabilities, we use high-throughput Thermofluor measurements for the 16 variants to extend analysis of the mutated residues to their impact on unliganded TrpRS stability. Aggregation of denatured proteins complicates thermodynamic interpretations of denaturation experiments. The free energy landscape of a liganded TrpRS complex, carried out for different purposes, closely matches the volume, helix content, and transition temperatures of Thermoflour and CD melting profiles. Regression analysis using the combinatorial design matrix accounts for >90% of the variance in T_m_s of both Thermofluor and CD melting profiles. We argue that the agreement of experimental melting temperatures with both computational free energy landscape and with Regression modeling means that experimental melting profiles can be used to analyze the thermodynamic impact of combinatorial mutations. Tertiary packing and aromatic stacking of Phenylalanine 37 exerts a dominant stabilizing effect on both native and molten globular states. The TrpRS Urzyme structure remains essentially intact at the highest temperatures explored by the simulations.

## INTRODUCTION

### TrpRS catalysis uses an escapement mechanism to transduce ATP hydrolysis free energy

Mechanistic studies of *B. stearothermophilus* tryptophanyl-tRNA synthetase (TrpRS) identified two, apparently contradictory, ways in which differential conformational stability affects its catalytic cycle ^1-3^. Catalytic assist by Mg^2+^ ion, on the one hand, depends entirely on its coupling to domain motions^3-5^. On the other hand, the sign of the free energy change for the catalytic conformational change is positive in the presence of the pyrophosphate product, PPi, and negative only in its absence ^6^. The former effect couples catalysis of NTP utilization tightly to domain motion; the latter ensures that domain motion requires PPi release, thereby reciprocally coupling the conformational change to the chemistry of ATP utilization. The two effects function analogously, one to each of the two blades of an escapement mechanism that converts pendulum oscillations into precisely-timed, unidirectional motions of the hands in a mechanical clock ^3^.

This novel escapement mechanism ^3^ ensures vectorial utilization of ATP ^7,8^, and is thus a potentially general model for energetic coupling in a broader range of enzymes that transduce NTP hydrolysis free energy ^9,10^. That deepened understanding of mechanical, biosynthetic, and informational coupling between domain motion and NTP hydrolysis provides a novel perspective on how biological processes transcend the second law of thermodynamics ^11,12^.

Experimental and computational data supporting this escapement mechanism were derived from a strategically selected set of combinatorial mutations of residues that mediate the shear developed during domain motion. That set positions us to probe in unprecedented detail the, as yet unknown, enabling links between differential conformational stability and allosteric function. The unusual array of structural TrpRS variants mean that such an effort requires precise, high-throughput characterization of structural melting. This paper establishes differential scanning fluorimetry (Thermofluor ^13,14^) as a suitable probe.

During tryptophan activation TrpRS proceeds through at least three distinct, metastable conformations: an Open ground state in which the Rossmann fold and anticodon-binding domains are rotated away from one another, separating the ATP and amino acid binding sites by about 6 Å ^15^; a Pre-transition (PreTS) state in which the domains are closed and in which the binding of Mg^2+^•ATP induces a 10 degree twist between them ^16^; and a Product state in which the twist is resolved, but the domains remain closed ^15,17,18^. We previously used binding affinity changes of two adenine nucleotide ligands to estimate that the twisted PreTS conformation increases conformational free energy by ∼3 kcal/mole ^19^. Moreover, removing ligands from MD simulations led either to rapid regression of the PreTS conformation to the open state, or to progression of a putative transition state analog complex to the product state ^20^. The closed, PreTS state is thus an excited conformational state formed before the chemical steps displacing the pyrophosphate group by the tryptophan carboxylate. Stabilization of the chemical transition state by TrpRS occurs transiently, during the untwisting motion ^1^.

Molecular dynamics simulations based on different crystal structures ^16^ show that the connecting peptide 1 (CP1) insertion in the TrpRS Rossmann fold and anticodon-binding (ABD) domains undergo anticorrelated motions, with CP1 and ABD moving as rigid bodies without internal core packing changes ^16,20,21^ during induced-fit and again during catalysis and tRNA acylation. These motions create shear that is mediated by alternative packing of seven residues (I4, F26, L29, Y33, C35, F37, I140) composing the D1 switch (Fig. 1). These residues are all contained within the TrpRS Urzyme, a highly active core structure conserved in all 10 Class I aminoacyl-tRNA synthetases (aaRS) ^22-25^. The centroid of the D1 switch, is ∼20 Å from the ATP α-phosphate.

**Fig. 1.**
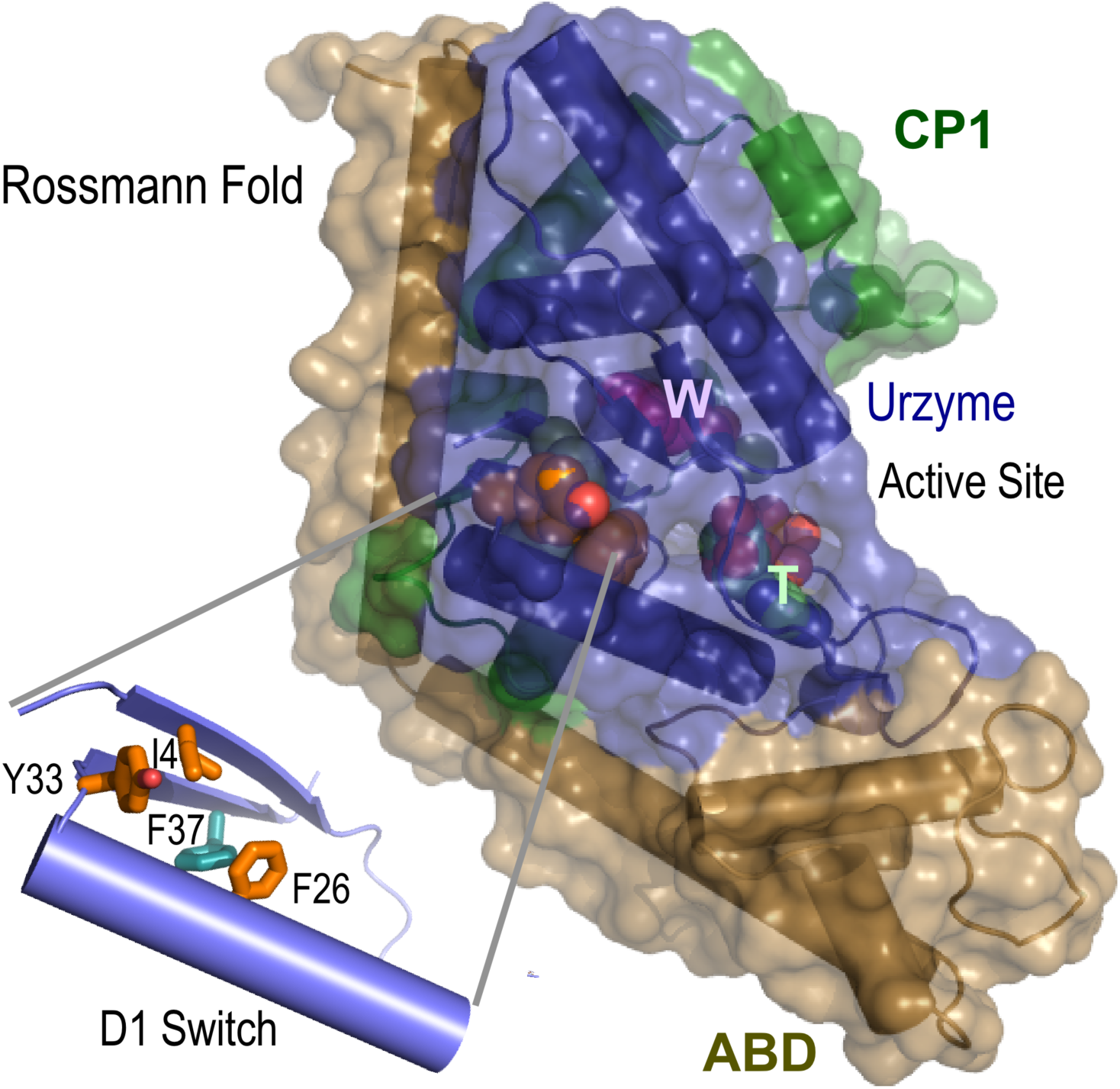
The D1 Switch mediates relative movements of two rigid bodies. The anticodon-binding domain (ABD; sand) and Connecting Peptide 1 (CP1; green) move essentially as rigid bodies relative to the Urzyme (blue) throughout the catalytic cycle. T and W indicate substrates Mg^2+^·ATP and tryptophan, respectively, in the active-site. Insert highlights the fact that the F26*Y33 interaction is local, as the residues are separated by only two α-helical turns, consistent with its major stabilizing effect on the α-helices. Stacking between F37 (teal) and F26 (orange) is essentially a tertiary interaction across the loop connecting the second β-strand to the first alpha helix, consistent with its dominant stabilizing role in the Molten globule states.

### Combinatorial mutants played a key role in identifying the escapement mechanism

Variants of four residues from the D1 switch were designed, using Rosetta ^3,20^, to minimize stability difference between the PreTS excited state and the preceding and following ground-state structures, in order to study how interactions between these residues impact catalysis. Fifteen combinatorial TrpRS mutants were constructed representing a full, 2^4^ factorial design including all possible perturbations of the four variants suggested by Rosetta. Multidimensional thermodynamic cycle analysis revealed that the five-way eregetic copuling of all four D1 residues with the active-site Mg^2+^ ion contributes –6.0 kcal/mole to transition-state stabilization during tryptophan activation ^3^ and ∼ –5.0 kcal/mole to the relative rates of tryptophan vs tyrosine activation ^26^.

The magnitudes of these highly cooperative, long-range catalytic effects, and the dynamic repacking associated with the untwisting catalytic domain motion ^1,16^ suggest that the D1 switch senses and communicates the domain configuration to the active-site Mg^2+^ ion, switching it to a catalytically active position only as the domains untwist. We validated that conclusion by showing that a modular thermodynamic cycle constructed from the full-length TrpRS, TrpRS Urzyme and intermediate constructs made by restoring to the Urzyme each of the two domains deleted in the Urzyme yields quantitatively the same coupling energy between the two domains ^27^. Mg^2+^-assisted catalysis thus arises entirely from domain motion ^3,26^.

More recently, minimum action path simulations implicated repacking of D1 switch residues in the conformational transition state ^6^. For each variant, trajectories furnished computational estimates of the conformational free energy difference between pre-transition state and products conformations, ΔG_conf_; the time to the conformational transition state, –ln(t_left_); and the conformational barrier height, U^‡^ ^28^. Remarkably, those variant-dependent computational estimates correlate closely with design matrix of mutant-site locations, and their impact on both k_cat_, and the single turnover rate, k_chem_ ^1,28^. The thermodynamic cycle relating structural, kinetic, and computational parameters establishes a unique consistency with the multi-dimensional thermodynamic cycles from which the escapement mechanism was first identified.

### Further progress requires precise, high throughput conformational stability analyses

Circumstantial evidence summarized in previous paragraphs suggests that the significant energetic coupling evident in the catalytic assist by Mg^2+^ and in amino acid specificity are emergent properties whose roots lie in differential conformational stabilities that result from changing interactions between the D1 switch residues. Those changes in conformational stability are, in turn, induced by the succession of bound ligands, as the chemical events proceed ^2^.

Differential scanning fluorimetry (Thermofluor; ^13,14^) provided preliminary evidence regarding how the four single D1 point mutants alter these relationships, relative to native TrpRS ^4^. Evidence on how the four side chains cooperate to produce catalytic effects, however, requires comparing native with all fifteen variants (see supplement to ref ^3^). Melting temperatures for the ensemble of mutant proteins would allow us to evaluate the intrinsic effects of each mutation, together with all six two-way, all four three-way, and the single four-way interaction.

Our goal is to establish a basis for pursuing two non-trivial longer-range questions: (i) Can we exploit these mutant proteins to correlate changes in the stability of all three conformations with the steady-state kinetic properties already measured ^3^? (ii) How do the interactions between the four mutated residues exert this highly cooperative effect? Can mechanistic patterns induced by the mutational perturbation inform the kinetic and thermodynamic descriptions of the catalytic cycle and eventually enhance our understanding of catalytic coupling? We use Thermofluor (differential scanning fluorimetry; ^13,29,30^) here together with CD to compare both probes of thermal melting for the TrpRS combinatorial variants. The two types of melting curve provide evidence that TrpRS unfolds via reversible formation of a molten globule intermediate state.

Quantitative comparisons between Thermofluor and CD melting curves allow us to identify statistically significant differences in how the D1 switch mutations alter the *T*_*m*_ values of native and intermediate molten globule states, relative to those of the fully denatured state defined by loss of ellipticity, θ_221_, arising from unfolding the α-helices. This analysis of unliganded TrpRS should help identify the intrinsic and higher-order structural interactions that allow the D1 switch residues to impact the stability of different conformational states along the structural reaction profile. Subsequent analysis of how the energetic couplings between these residues change in the presence of ligands representing the structural reaction profile can then be related to both experimental and computational parameters derived for the entire ensemble of variants.

### Aggregation complicates thermodynamic interpretations of denaturation experiments

A requisite for attributing thermodynamic significance to denaturation experiments is that the melting curves be reversible. Many proteins, however, fail to meet this exacting criterion, because their unfolded configurations aggregate to varying degrees at the concentrations necessary for high signal to noise measurements. TrpRS exhibits this behavior (Supplementary §1).

We adduce two complementary kinds of evidence that the underlying process is reversible, justifying thermodynamic interpretations. First, the structural and heat capacity changes for thermal unfolding of the TrpRS monomer in a computational free energy surface (that is, by definition reversible at infinite dilution) closely resemble those of the experimental transitions. Second, the combinatorial design matrix implies simultaneous equations whose least squares solutions for the vectors {T_m_,_obs_φ_i_} and {T_m,obs,_θ_1_} explain > 90 % of the variance in both observed melting temperatures, suggesting that errors arising from aggregation have only marginal impact on differences between T_m_’s for the ensemble of structural variants.

## METHODS

### Expression and purification

His_6_-N-terminally tagged and untagged WT TrpRS and mutant proteins were expressed in *E. coli* BL21(DE3)pLysS, with kanamycin and chloramphenicol. Cells were resuspended in 20mM HEPES pH 7.6, 0.3 M NaCl_2_, 10 mM BME, 30 mM imidazole (2.5 mL/g wet weight). Cell pastes were sonicated and cleared by centrifugation at 15000 rpm for 30 minutes at 4 °C. The cleared lysate was mixed with Ni beads (Thermo Scientific HisPur^Tm^ Ni-NTA Resin) at approximately 2-3 mL per liter of lysate and the suspension passed into a self-flowing glass column. After washing with 20 column volumes His-tagged protein were eluted with 20 mM HEPES pH 7.6, 0.3 M NaCl_2_, 10 mM BME, 0.3 M Imidazole. Imidazole concentrations were reduced by dilution to 30 mM before incubating overnight with TEV protease at a ratio of 1:10. Purified, cleaved proteins were concentrated using an Amicon PM10 Ultra membrane and stored at −20°C in 50% Glycerol. Roughly 30% of the experiments were performed on both tagged and untagged proteins, without significant differences. We report here only results using tagged and cleaved proteins.

### Combinatorial mutagenesis

Rosetta ^31^ was used to identify mutations to four of seven non-polar residues in the D1 switch ^20^, that participate in a broadly conserved ^23^ tertiary packing motif in the first β-α-β crossover connection of the Rossmann fold. Mutations I4V, F26L, Y33F, and F37I, were selected to minimize the increased conformational energy of unliganded PreTS TrpRS ^15^, relative to the Open ground ^32^ and Product ^17,33^ states identified by X-ray crystallography.

### Thermofluor Stability Assays

Differential scanning fluorimetry (Thermofluor; ^13,29,30^) was developed as a high-throughput method to compare ligand affinity from the elevation of melting temperatures. Thermofluor is especially useful for our purposes, because it can be performed in 384-well microtiter plates. Rapid, highly replicated analysis of many combinations of protein mutational variants and ligands affords experimental access to questions about higher-order interactions between conformational stability and ligand affinity.

A 6:5000 dilution of Sypro Orange dye into 20 mM HEPES, 50 mM NaCl_2_ (pH 7.0) buffer was mixed with 20 μl of a 6 μMol enzyme solution, with 20 mM MgCl_2_. No ligands were added. Each sample was replicated 6 times on the 384-well format plate, which was heated from 25° C to 95° C in 0.4 degree steps lasting 8.7 sec. Fluorescence measurements were done with an ABI 7900HTFast Real-Time PCR instrument and data were analyzed either with SigmaPlot or with MATLAB programs described further below. One-off melting curves using ANS (1-Anilino-8-naphthalene sulfonate) were performed using a SPEX Fluorolog-3 spectrofluorometer. Identical, six-fold replicated measurements were performed mutant proteins purified on two different occasions, substantially reducing the overall variance of estimated melting temperatures. Subsequent experiments were performed in this manner with 1 mM MgCl_2_, 1mM MnCl_2_, and without added metals.

### Circular dichroism (CD)

Circular dichroism measurements were done on an Applied Photophysics Chirascan Plus steady-state Circular Dichroism instrument in the UNC Macinfac facility. We measured the ellipticity of 8 μM of each protein in 200 mM Sodium Phosphate Buffer (pH 7.3) without added metals from 25° C to 90° C at 221nm (θ_221_) and similarly at 270 nm (θ_270_).

### Data reduction and parameter estimation

Both Thermofluor and CD melting curves were processed using Matlab® (Mathworks ^34^. The software was built as a pipeline of .m-files that reduce and analyze the data, including thermodynamic characterization and presentation of statistics. The pipeline consists of three parts:

1. Reading data from high-throughput RT PCR files and transforming them into a matrix with four columns: i) well number; ii) an index representing the mutation (all mutants were assigned a number so the wild type was indexed by 0, I4V by 1, and so on, according to the sequence number of the mutation(s)); and finally the data, iii) temperature and iv) fluorescence reading.
2. Fitting the data (both Thermofluor and CD)
3. Determination of *T*_*m*_ by the “ratio” method (see Supplementary §S2).

Outputs are: error log, parameter file, an output Excel data file with fitting and simulation parameters, and an Excel file where replicates are averaged and recorded with standard deviations and errors.

The software determines the temperatures at the low and high levels of the sigmoidal part of the experimental curve. If these cannot be found, the user is prompted to input them manually. These positions determine the intervals for the initial, and final linear baselines and the sigmoidal segment (Supplementary Eqs. (S3)). These intervals then are used for the initial parameter estimation in the “Model” scenario and to extrapolate the initial and final parts to utilize the “Ratio” method. An additional module estimates T_m_ using a thermodynamic module. The program allows for repeated analysis and/or re-initialization for problematic data sets if needed. Matlab codes are included in Supplementary §9.

### Replica Exchange DMD simulations

The replica exchange algorithm efficiently samples the temperature-dependent conformational free energy landscape of a macromolecule by simulating replicas of the equilibrium state at different temperatures and, at predefined time points, exchanging the structures at different temperatures if the difference in their energies is within a threshold. We implemented the replica exchange algorithm using the REX/DMD suite ^35-38^ that uses Discrete Molecular Dynamics ^36,38-40^ to simulate the replicas at 24 temperatures ranging from ∼175K to ∼405K separated by a constant interval. DMD approximates atomic interactions by multistep square well potentials ^35,41^ and uses an Andersen Thermostat to regulate the temperature of the system while Lazaridis-Karplus implicit solvation model ^42^) accounts for the solvation energy. Simulations were set up with the PreTS state structure, 1MAU, which appears to be the least stable of the crystal structures.

We excised the terminal amino acid (R328) from the structures as it is not observed in most crystal structures. Also, since DMD force field does not include parameters for Mg^2+^, we replaced it with Zn^2+^. We do not expect these changes to affect the dynamics of the system in a significant manner. To retain the ligands in the binding pocket a harmonic constraint (well width = ±1Å) was applied between the atoms of the ligands and the binding pocket (within 3.5 Å of each other). Another weak harmonic constraint (well width = ±2Å) is applied to atoms which uniquely form native contacts (8Å), in order to facilitate the exploration of the conformational space adjacent to the low energy equilibrium state. The simulations at each temperature were run for a total duration of 3 million steps (∼150 ns). Omitting the time to equilibrate (initial 500,000 steps), snapshots were generated every 1000 steps for a total of 2500 snapshots which were then used in all the analyses.

### Regression modeling ^3,4^

Horovitz and Fersht outlined how multiple mutant cycles give evidence for energetic coupling between sites by evaluating all intersecting two-way thermodynamic cycles ^43^. Misconceptions nonetheless persist arising from confusion between the actual experimental observations for individual multiple mutants themselves, and the coupling energies they imply. Energetic coupling between residues depends on how the effect of a particular mutation changes in different contexts (wild-type, mutations at other sites, etc.) in which it is observed. For that reason, the entire set of combinatorial mutants must be analyzed as an ensemble. Comparison between a quadruple mutant and WT protein may suggest only a small interaction when, in fact, the overall coupling can be quite significant. Our own work ^3,4^ and that of others ^44^ provide examples of how higher-order coupling emerges from multi-mutant thermodynamic cycles.

The computational tedium of working out individual two-way thermodynamic cycles can be minimized by using regression modeling to solve the simultaneous equations implicit for a high-order thermodynamic cycle to estimate contributions to thermal stability from intrinsic mutational effects and higher-order interactions jointly. Adjusting coefficients to minimize the total squared residual is straightforward in many statistics programs, e.g. JMP ^45^, and leads simultaneously to maximum-likelihood estimates of the parameters and their standard errors. Such calculations are otherwise equivalent to estimating coupling effects individually, as described ^43^.

Regression methods have two additional non-trivial advantages: (i) The squared correlation coefficient between 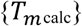 and 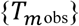, R^2^, is equal to the fraction of the variance in 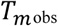 that can be explained by the model. Values close to 1.0 favor causal relationships. Moreover, high precision of the data collection is key to a conclusive interpretation of melting behavior, because the maximum R^2^ that can expected for a given model, in turn, decreases linearly with the square of the relative error in the experimental data to which it is fitted ^46^. This internal consistency of a regression model furnishes a metric for the extent to which the model may have omitted important causative variables. (ii) Ratios of model coefficients to their standard errors provide Student t-values, whose probabilities under the null hypothesis are known, giving statistical significance for each estimate. The relative size of the error vector, **ε**, the implied statistical significance of the Student t-tests, and the R^2^ value associated with regression models thus constitute an interrelated set of metrics for the degree to which the melting temperatures have thermodynamic meaning.

## RESULTS AND DISCUSSION

We compare melting behavior for all unliganded variants observed by Thermofluor and CD at 221 nm (θ_221_). These data suggest successive, linked, quasi-two-state processes leading ultimately to the loss of secondary structure. We consider requirements for equilibrium measurements and reversibility, using data from related controls in Supplementary §1, §2. Evidence from REX/DMD temperature-dependent free energy surfaces also shows that TrpRS unfolds in two stages with much the same behavior as that seen in the experimental melting curves. Thus, monomeric TrpRS denatures reversibly, consistent with thermodynamic interpretations of the experimental melting curves. Linear regression models based on the combinatorial design matrix consistently attribute > 90 % of the variance in {*T*_*m*_} for both processes quantitatively to mutational effects. We discuss how that linearity provides orthogonal and complementary evidence that the experimental melting curves contain thermodynamically meaningful information. Finally, the computational melting profile suggests that the TrpRS Urzyme remains largely intact at 94 C.

### TrpRS denatures via an intermediate, molten globular state

Experimental Thermofluor and CD melting curves are shown in Fig. 2 for all variants. Mean *T*_*m*_ values for both processes are given in Supplementary Table SI. Variances from fitting the CD melting curves are comparable to those obtained by averaging six-fold replication in the 384-well plate and from repetition of the entire plates on two different dates with different TrpRS preparations. The mean increase for all mutants between 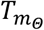 vs 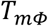 is 8.5° ± 0.8°.

**Fig. 2.**
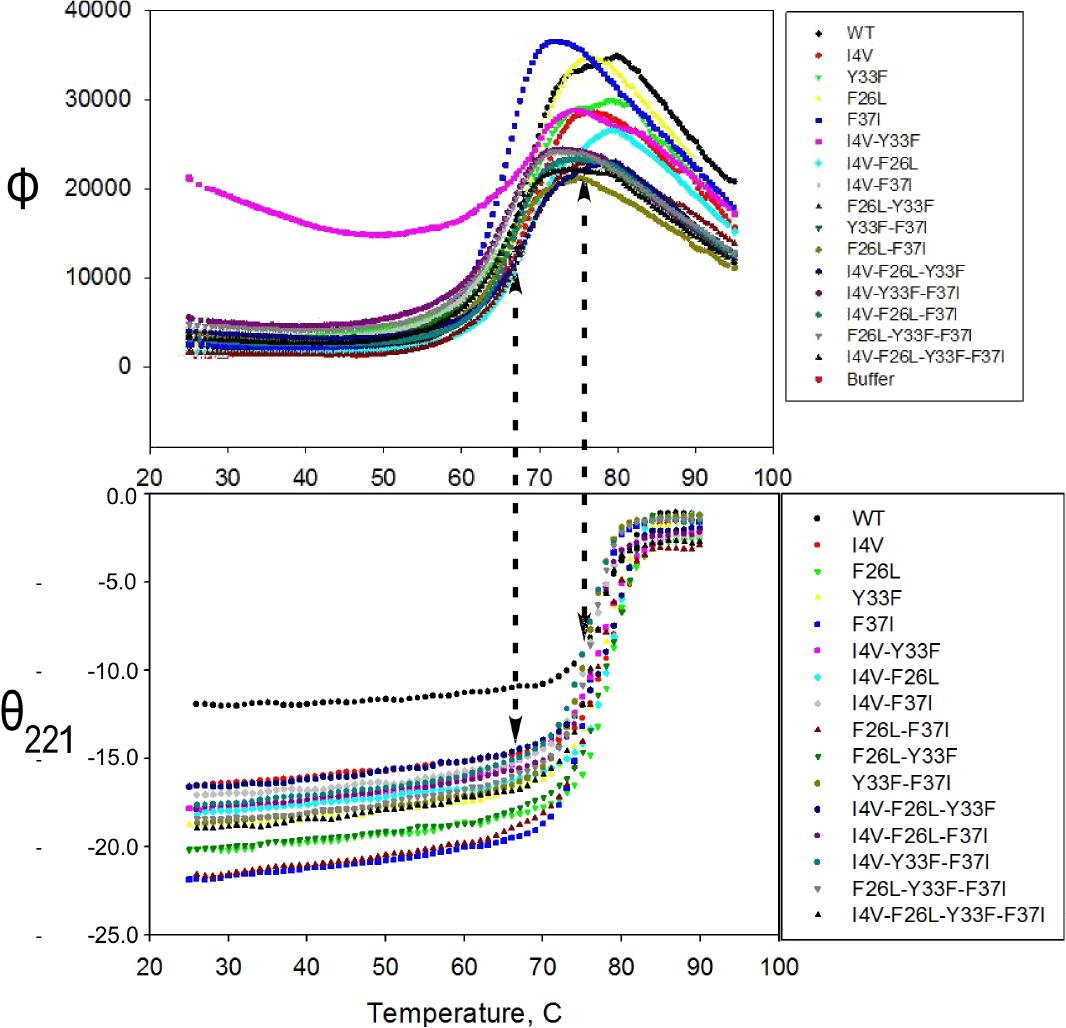
Raw data for all thermal melting processes. Vertical dashed arrows align the midpoints of the two transitions with corresponding progress along the other process. Note that the melting of α-helices does not begin until the midpoint of the molten globular transition.

#### Two-state thermodynamic modeling, model-free estimation, and computational free energy simulations agree on successive TrpRS melting temperatures

*T*_*m*_ estimates from fitting to a conventional two-state model ^48^ are compared in Supplementary Table SI with those estimated by the ratio method (Fig. S3).

#### Thermofluor, θ_221_, and simulations detect physically different changes

Ellipticity at 221 nm arises from α-helical secondary structures, whereas Thermofluor detects partial breakdown of native-like non-polar side chain packing. The latter process is related to the loss of ellipticity at 270 nm (Fig. S1C), which arises from persistent asymmetries in the relative positions of aromatic sidechains ^49^.

#### Controls ensure that the schedule of increasing temperatures allowed sufficient time for equilibration and test the reversibility of TrpRS denaturation

These controls are described in Supplementary §1 and illustrated Fig. S1. Although the time interval between temperature changes should allow sufficient time for equilibration at each temperature (Fig. S1B), there is evidence in Fig. S1C, D for partial losses owing to aggregation of denaturated TrpRS. These losses contradict the assumption of reversibility.

#### Protein concentration effects

That TrpRS is a functional dimer introduces another complicating factor because melting curves will depend to some extent on protein concentrations related to the dimer dissociation constant, which must also interact with monomer denaturation. The dimer dissociation constant measured directly from monomer/dimer distributions using atomic force microscopy is K_D_ = 5 nM in a similar buffer (Lewis, S.; Na, M.; and Erie, D.; Carter, CW Jr; UNC unpublished). As we measure melting at much higher concentrations (ie. 6-8 μM), we expect this effect to be minimal as well. Supplementary §2 discusses experimental evidence that *T*_*m*_,s measured over a four-fold concentration range differ by <1 degree Celsius (Fig. S2).

#### State probabilities can be estimated as a function of temperature as described in supplementary §6 (Fig. S4)

The correspondence implies that the second melting process requires prior loss of native-like core packing, according to a three-state scheme. Cremades and Sancho describe similar results with flavodoxin ^50^. Melting transitions detected by Thermofluor and θ_221_ (Fig. 2) show that α-helices do not begin to melt until the molten globule concentration reaches a maximum, and are completely melted only when the molten globule concentration is exhausted, as in the REX/DMD simulations (Fig. 4). Ptitsyn and co-workers ^49^ reported a similar observation for α-lactoglobulin. All mutants show a pattern (Supplemental Fig. S4) in which (Supplemental Fig. S3: (i) P_2max_ is close to 1.0; (ii) the probabilities P_1_ and P_3_ at P_2max_ are close to 0).

**Fig. 3.**
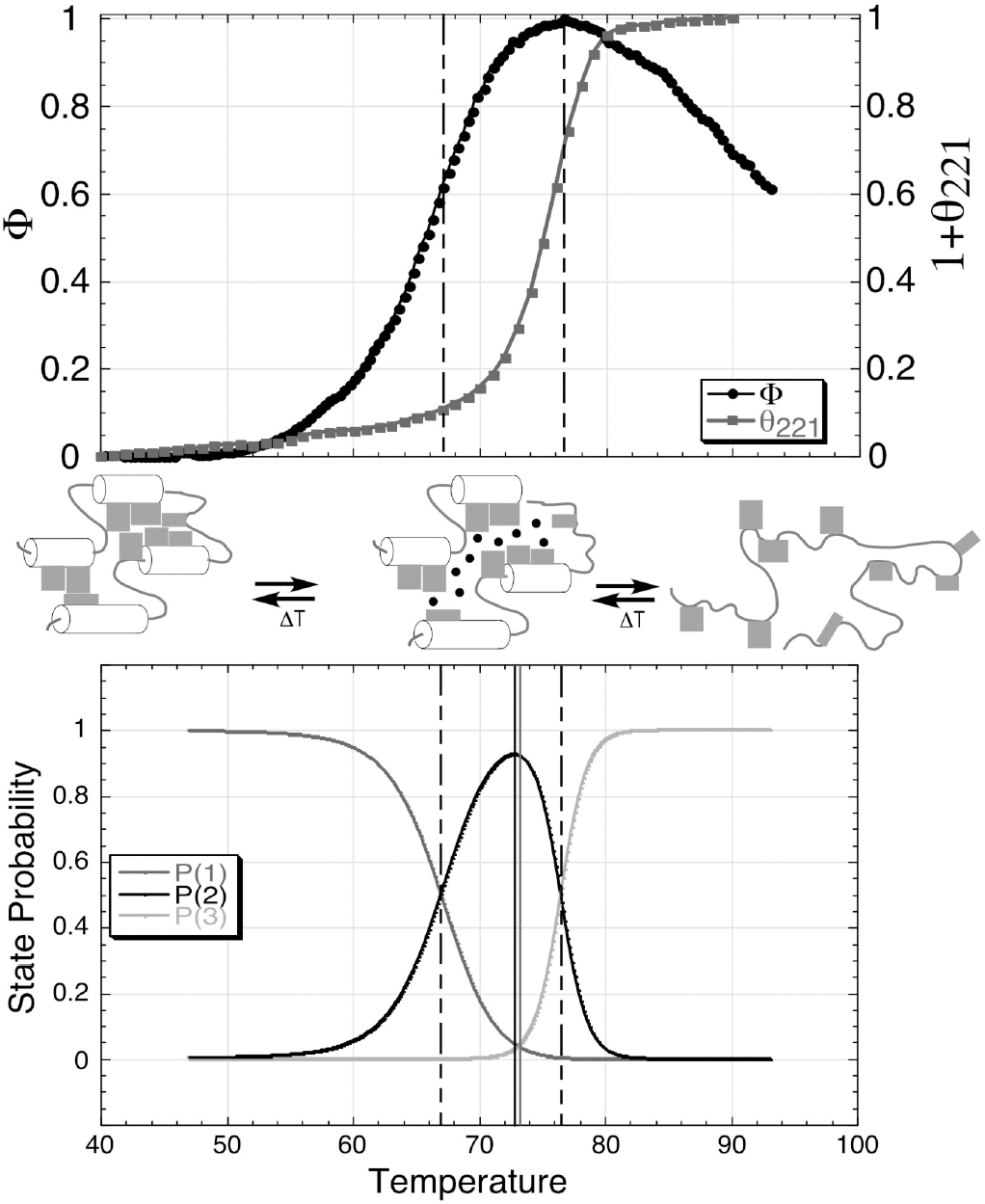
Quantitative metrics supporting a three-state model for native TrpRS denaturation. A. Normalized experimental Thermofluor (circles) and CD (triangles) melting curves. The same temperature scale in degrees celsius applies to all three panels. Dominant structural ensembles below *T*_*m*_ (Thermofluor) resemble the native state, S1. Between *T*_*m*_(Thermofluor) and *T*_*m*_(CD) they resemble molten globule, S2, and above *T*_*m*_ (θ_221_) they are characterized by loss of α-helices detected by CD. Vertical lines represent melting points detected by the two data sets. B. Probabilities (P_1_, P_2_, P_3_) of the three states represented in A. The temperatures and amplitudes of P2 and the intersection of P_1_=P_3_ are not constrained by Eqs. (9) and hence define quantitative metrics of the validity of the three-state approximation.

**Fig. 4.**
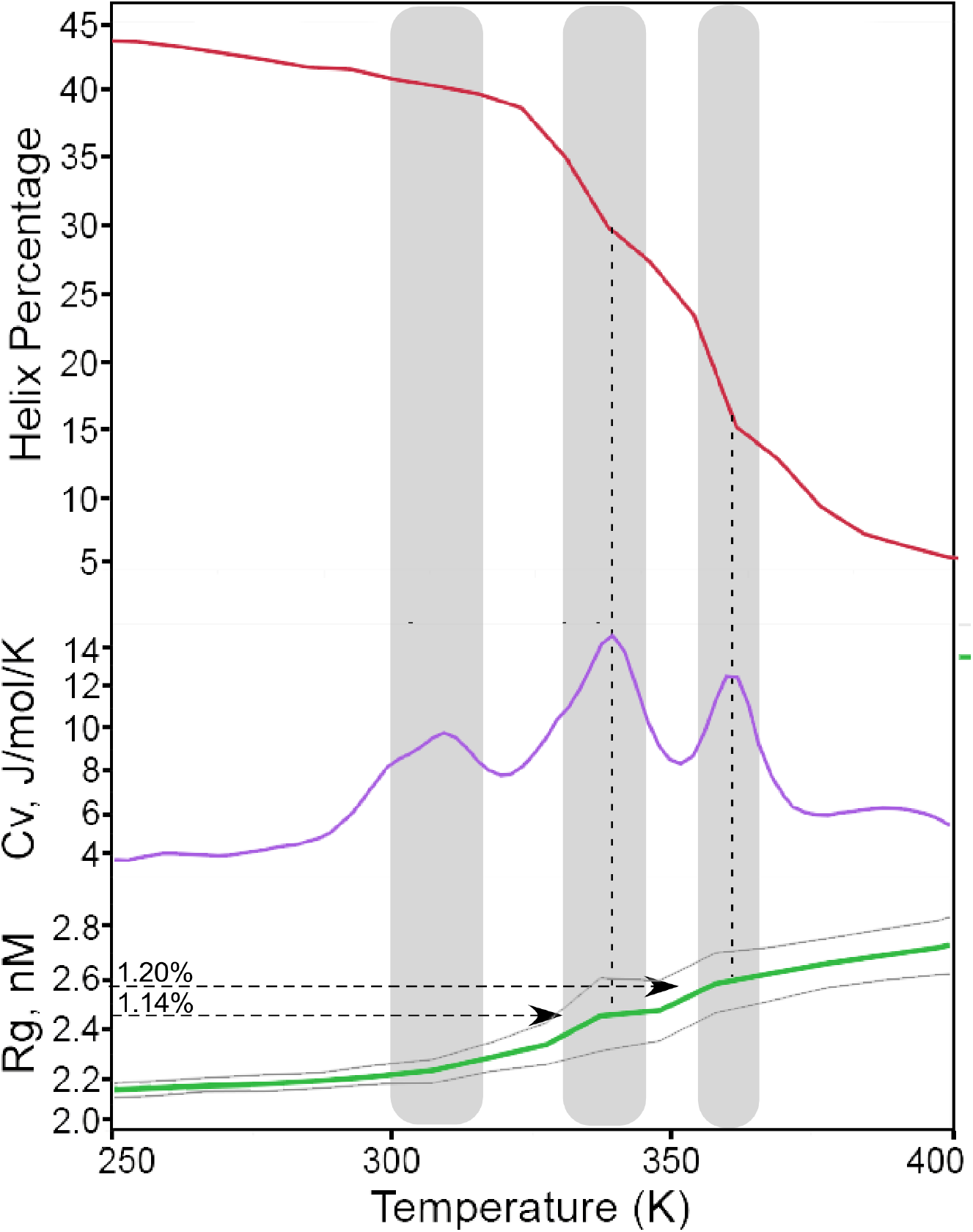
Computational traces of TrpRS thermal denaturation. The grey shading highlights progressive maxima in the specific heat capacity, C_v_, (blue curve). The two, principal, symmetric peaks occur at temperatures corresponding to those measured by Thermofluor (67 C vs 69 C) and ellipticity (87 C vs 80 C). Both transitions also correspond to increases in volume, as indicated by the radius of gyration (R_g_, green curve, bounded by its uncertainties in grey). The 14% increase in R_g_ in the first transition corresponds to that (15-20%) expected from formation of a molten globule ^54^. Similarly, the loss in helical content (red curve) occurs in two stages, the more dramatic of which coincides with the second transition. Vertical dashed lines align discontinuities in both structural parameters with the stationary points of the C_v_ trace.

Two different probes identify transitions between states at two different temperatures. They therefore rule out two-state denaturation and strongly imply formation of an intermediate, molten globular state ^47^. TrpRS thermal denaturation therefore appears to be a three-state process involving two successive transitions, each of which is two-state to good approximation, and for which the intermediate molten globule state is the reactant in the second. We show in a subsequent section that perturbations induced by specific mutations on the two stability measurements have similar, though not identical effects on molten globule formation and loss of α-helices, as detected by the respective structural probes.

Our conclusion that TrpRS denatures in two stages, the first of which is essentially molten globular in nature, implies that unfolding—a first-order process—occurs at a rate proportional to the concentration of reactant. The succession of states in the thermal structural profile is such that the unfolding of α-helices begins once the concentration of molten globules reaches roughly 50% of the total protein concentration. Stabilization or (less likely) destabilization of the molten globular state by dye binding would change the concentration of the reactant for the final unfolding of the helices in a regime where those changes occur most rapidly—i.e., where sypro orange fluorescence is changing most rapidly with temperature. Any change in *T*_*m*_φ, in turn, would affect the concentration of molecules that give rise to the final unfolding process, and be expected to induce changes in the earliest part of the CD melting curve. The fact that the CD melting curves with and without sypro orange cannot be distinguished over any of their sigmoidal shape thus does imply that the dye has no detectable influence on the formation of the molten globular state. Thus, although Sypro Orange may stabilize the molten globule very weakly, the effect is small enough to ignore for our purposes.

### A reversible REX/DMD free energy landscape reproduces transitions and structural changes associated with Thermofluor, CD melting curves for the TrpRS monomer

#### A computational temperature-dependent free energy surface of a liganded TrpRS closely resembles the experimental denaturation profile of unliganded TrRS

To validate the thermodynamic interpretations of TrpRS melting behavior, we take advantage here of detailed correspondences between the experimental melting curves (Fig. 2) and a computational temperature-dependent conformational free energy surface of the liganded, wild-type TrpRS monomer PreTS complex, mapped previously using Replica Exchange Discrete Molecular Dynamics (REX/DMD; ^51-53^) simulations to confirm the free energy landscape associated with computational transition states ^1,6,28^. That surface shows transitional events at three temperatures comparable to those observed experimentally. These features are reversible, by definition of the free energy surface. Although that temperature-dependent free energy surface was performed for a liganded, monomeric species, it has the advantages of single molecule studies in which aggregation between molecules cannot interfere with reversibility ^53^.

Computational estimates of the radius of gyration (R_g_), heat capacity, and helical content of the liganded TrpRS monomer as functions of temperature are shown in Fig. 3. All three properties exhibit excellent correspondences between the computational free energy simulations and the Thermofluor and CD traces. The computational radius of gyration increased in two steps with increasing temperature that correspond to the second and third peaks in specific heat capacity plots. The average radius of gyration increases by 14% during the first transition which corresponds to the increase expected for a molten globule transition (15-20%) ^54^. At the 67 C transition temperature for the second peak of the specific heat capacity curve, 70% of the helices present in the native protein remain intact. At 87 C, the third transition temperature, only 28% of the native helices remain intact. The computational data therefore support the three-state denaturation mechanism implied by the experimental melting, in which reversible partial loss of tertiary structure precedes loss of secondary structure.

From the energies of structures generated at different temperatures the specific heat capacity, *C*_*v*_, can be computed as a function of temperature using the WHAM algorithm ^55^. *C*_*v*_ for PreTS exhibits three peaks as a function of temperature, two of which are associated with significant changes in structural parameters. The second peak at 67 C corresponds to 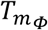 and the third peak was at 87 C corresponds to 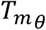. These computational estimates, understandably, differ somewhat from those obtained experimentally. The qualitative similarity is, nonetheless, remarkable.

Each heat capacity peak corresponds to a different transition because maxima in specific heat capacity are stationary points where the protein absorbs and then loses excess heat in the system as it undergoes conformation changes, so the system temperature remains constant. Curiously, discontinuities in the R_*g*_ and helix content coincide with local maxima in the *C*_*v*_ trace. Evidence corresponding to the first peak at 36 C in the computational also appears at ∼ 60 C in normalized plots of both experimental fluorescence and ellipticity traces in Figs. 2 and 3. This peak in the *C*_*v*_ trace has a smaller amplitude and involves minimal changes in structural metrics. It is evident mainly from the that it initiates the increasing variance of R_g_, which occurs almost exclusively with the formation of the molten globule. As 60 C coincides with the optimal growth temperature for *Bacillus stearothermophilus*, it is possible that this local maximum in *C*_*v*_ has biological significance. However, unpublished temperature dependent activity curves for native TrpRS (V. Weinreb) are not informative on this point.

#### The correspondence between experimental Thermofluor and CD melting curves (Fig. 2) and computational occurrences of these two transition events (Fig.4) is thus remarkably detailed

That agreement implies that, although complete reversibility cannot be demonstrated for solution ensembles of dimeric TrpRS molecules, the Thermofluor and CD melting curves nevertheless confirm the behavior observed in the reversible single molecule simulations, supporting thermodynamic interpretation.

### All Tm values are related linearly to the pattern of mutational sites

#### Multiple regression analysis demonstrates linearity of the impacts of mutation on apparent stability

Calvin, Hermans, and Scheraga ^56^ derived a linear relationship between the proportional change in melting temperature and the incremental free energy change induced by perturbing a reversible equilibrium between two differently folded states of a macromolecule (eqn. 1):

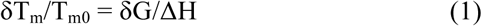

where T_m0_ is the melting temperature of a reference, or WT TrpRS, and ΔH is the overall enthalpy change of its structural transition. The matrix of combinatorial variants represents just such perturbations in a balanced, factorial design. Proportionality of δT_m_ to free energy implies, in turn, that thermodynamically meaningful values should be additive. The vector of 16 melting temperatures for each transition thus provides an implicit test of that relationship. The barcode in Fig. 5(A) represents the design matrix for the combinatorial mutagenesis, and that design matrix implies a set of simultaneous equations for T_m calc,i_ (eqn. 2):

**Fig. 5.**
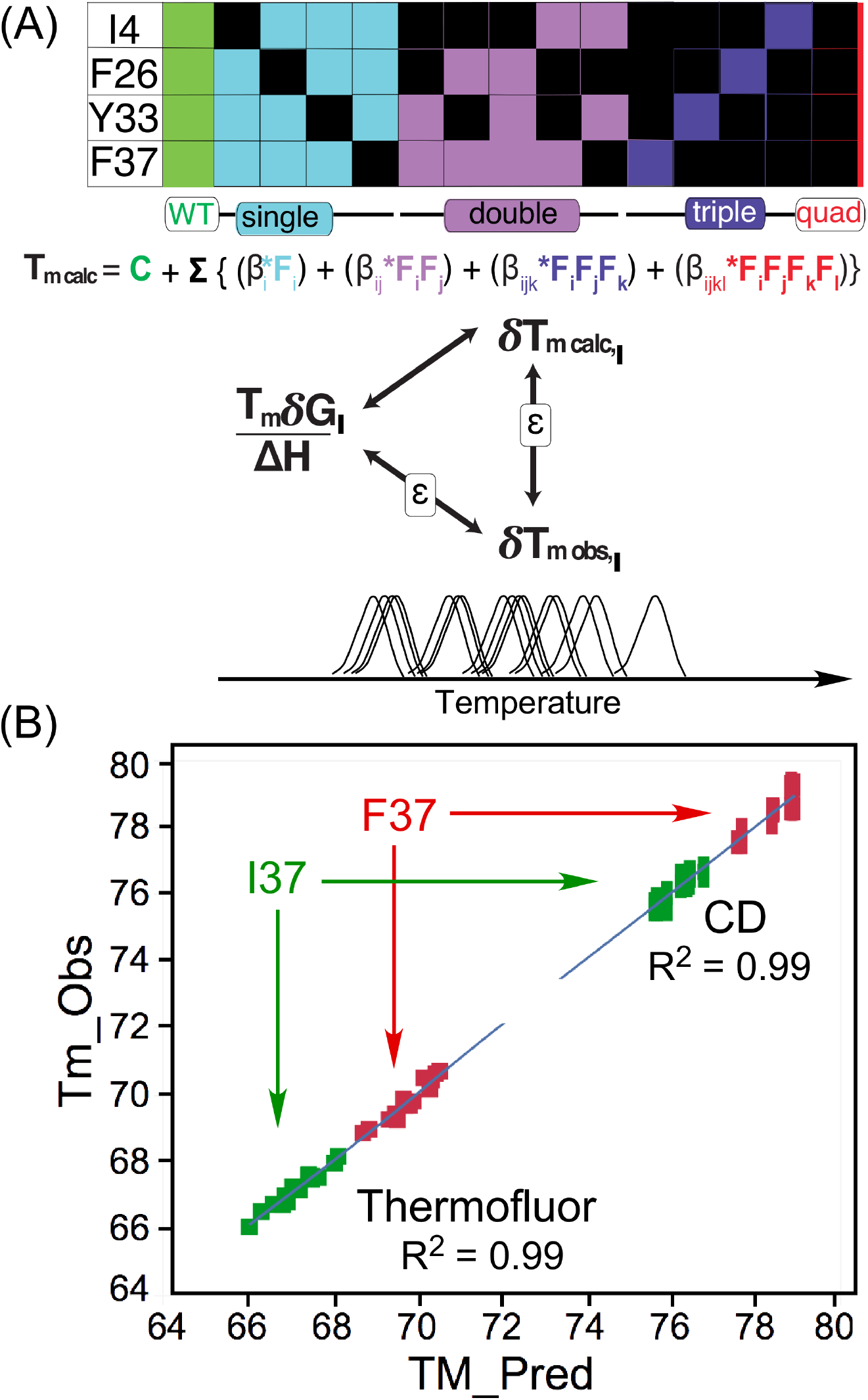
Linear self-consistency of T_m_ values calculated from regression models. (A) Thermodynamic cycle illustrating the argument that the extent to which calculated and observed {δT_m,i_} values agree is a measure of how closely the measured values approach proportionality to incremental free energies,{δG_i_}. The colored barcode in the upper part of the diagram represents the factorial design of combinatorial mutations and differentiates the four types of predictors in the simultaneous equations of the regression model. Observed T_m_ values are suggested schematically by the first derivatives of melting curves in Fig. 3. (B) Calculated T_m_ values for wild type and each of 15 combinatorial D1 switch mutants agree closely with observed values for both melting transitions, validating the regression models and supporting the argument illustrated in (A). Note that both 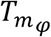 and 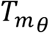 values are clustered, according to whether residue 37 is F or I. Although 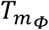 and 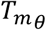 have different β coefficients, Table I, the two sets of modeled melting temperatures are nearly co-linear.

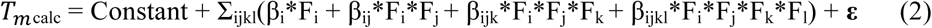

where βs are coefficients in degrees, {F_i,ij,ijk,ijkl_} = {0 or 1} are binary elements of the design matrix describing the presence or absence of each mutation as illustrated by the barcode, and 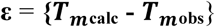 is a difference vector. By (1), **{δT**_**m calc**,**i**_**}** is proportional to **{δG**_**i**_**}** if all variants have the same enthalpy change.

The smaller the difference vector, **ε**, the better the approximation **{δT**_**m calc**,**i**_**}** is to **{δT**_**m obs**,**i**_**}**, and the better the approximation **{δT**_**m obs**,**i**_**}** is to **{δG**_**i**_**}.** Agreement between observed and calculated T_m_ values thus corroborates the linear relationship (1). Evidence of linearity in the Thermofluor measurements in Fig. 5(B) is derived from replicated measurements from two different TrpRS purifications made three months apart and in the presence of 20 mM MgCl_2_. The effects of divalent metal ions on Thermofluor melting curves are discussed further in Supplementary §S7 and illustrated in Fig. S5(A). Mg^2+^ ion lowers the Thermofluor melting temperature of all variants by ∼2.5 C.

The high R^2^, 0.99, of both Thermofluor and CD multiple regression models simultaneously imply that: (i) melting transitions for different mutants have similar overall enthalpy changes; (ii) ensembles of microstates that accumulate maximally at the transition temperatures are approximately at equilibrium, despite the observed aggregation (Supplementary §1); (iii) differences in their free energies arise primarily from entropic differences induced by the mutations. The evident linearity and vectorial nature of the combinatorial mutagenesis therefore stengthen the evidence from the free energy surface (Fig. 4) that TrpRS melting curves have meaningful thermodynamic interpretations.

#### Multi-mutant thermodynamic cycles yield structural insight into the molten globule state

The regression model for a 2^4^ factorial design ^57^ has 16 independent parameters, including the constant term. Replicated measurements therefore permit evaluation of all coefficients, with numerous degrees of freedom. The sixteen variants balance comparisons of mutant behaviors. The intrinsic effect of each mutation is averaged over eight contexts, that of each double mutant over four contexts, and each triple mutant over two different contexts (wild-type and mutant in the remaining residue). Regression modeling uses this averaging to enhance the precision and sensitivity of mutational analysis, quantitation, and significance testing of high-order interactions (see supplement to ref ^3^).

Factorial mutagenesis of the D1 switch region affords the opportunity to attribute stability changes in the two successive melting steps quantitatively to effects and cooperative interactions between specific residues. Fig. 5(B) shows the high correlation between *T*_*m*_ values calculated from the regression models and those observed experimentally. The regression models are summarized in Table 1. *T*_*m*_ values estimated from both Thermofluor and CD melting curves and sorted in ascending order are shown in Fig. 6(A), (B). The eight variants with wild-type F37 (Black histograms in Fig. 6(A), (B) are all more stable against loss of both native and molten globular structure than are the remaining variants with the mutant I37. All but one of the remaining mutant variants, by contrast, are more stable than wild-type TrpRS in both melting processes.

**Table 1.** Regression coefficients and Student test P values of predictive models illustrated in Fig. 5 for 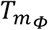 and 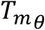. R^2^ values for the models are 0.99, 0.99, and root mean squared errors are 0.21 and 0.28 degrees, respectively.

| Predictor | $\beta_{\phi}$ | P$_{\phi}$ | $\beta_{\theta}$ | P$_{\theta}$ |
| --- | --- | --- | --- | --- |
| I4 | -0.28 | 0.196 | -0.06 | 0.783 |
| F26 | -0.06 | 0.772 | -0.76 | 0.001 |
| Y33 | 0.55 | 0.020 | 0.02 | 0.942 |
| F37 | 2.79 | <.0001 | 2.38 | <.0001 |
| I4*F26 | 0.98 | 0.005 |  |  |
| I4*Y33 | -0.09 | 0.755 | 0.38 | 0.193 |
| I4*F37 | -0.68 | 0.037 | 0.17 | 0.557 |
| F26*Y33 | -1.05 | 0.003 | 0.22 | 0.457 |
| F26*F37 | 0.10 | 0.748 | -0.50 | 0.096 |
| Y33*F37 | -0.69 | 0.033 | 0.05 | 0.883 |
| I4*F26*Y33 | -0.88 | 0.053 |  |  |
| I4*F26*F37 | 0.17 | 0.691 |  |  |
| I4*Y33*F37 | 1.43 | 0.004 | -0.95 | 0.028 |
| F26*Y33*F37 | 1.30 | 0.007 | 1.00 | 0.022 |
| I4*F26*Y33*F37 | -1.60 | 0.017 |  |  |

**Fig. 6.**
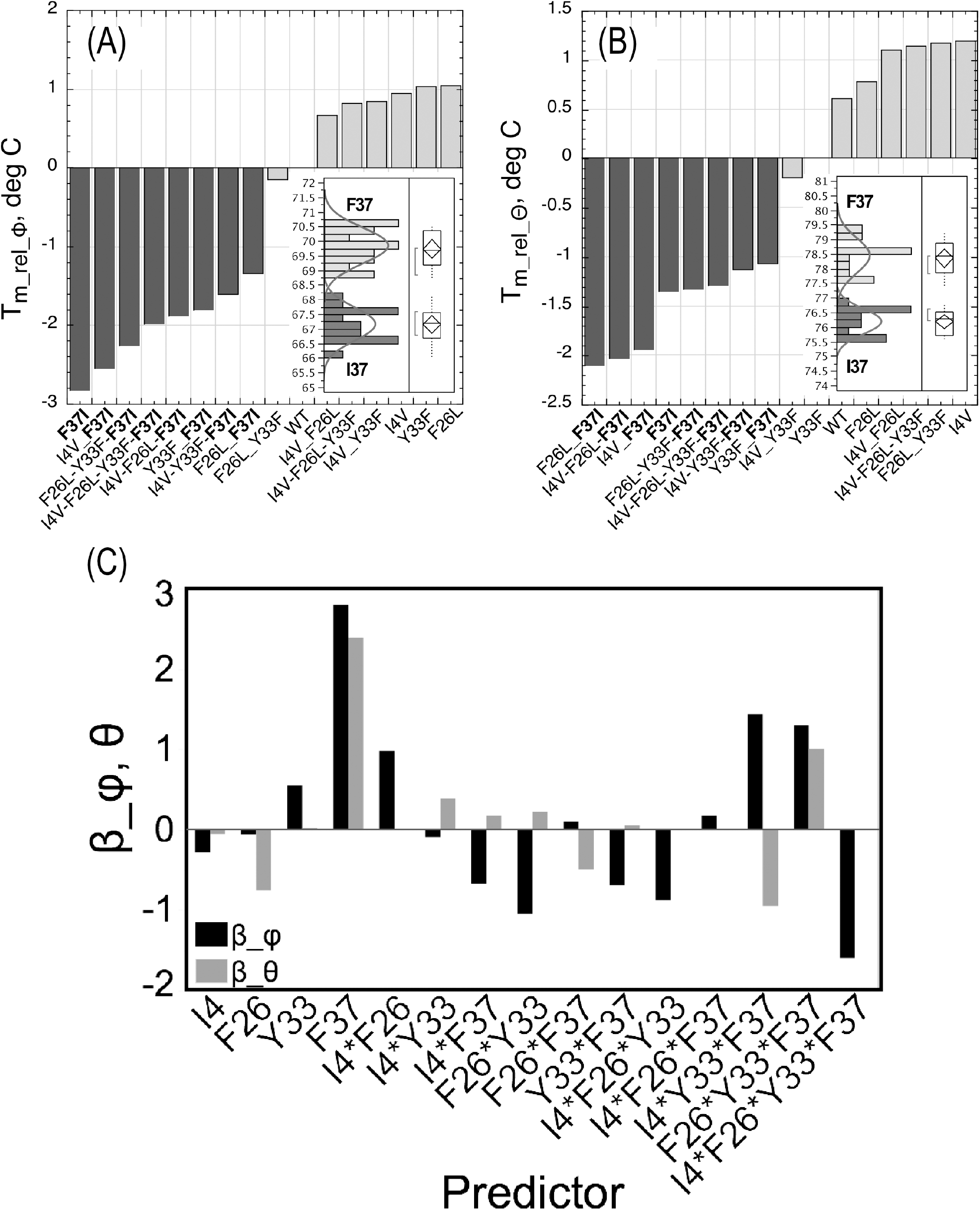
Differential effects of the 15 mutant proteins on the relative increase in 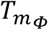 and 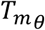. Similar factors contribute to melting temperature variations for molten globule formation (A) and loss of α-helices (B) in unliganded TrpRS D1 switch mutants. Wild Type F37 residue interactions are dominant and independently stabilize both MG and secondary structure (Prepared with Kaleidagraph ^72^). (C). Comparison of major contributions to the relative stability of TrpRS α-helices and molten globular states. Coefficients of regression models for 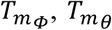, prepared using the program, JMP (*25*). Quantitative estimates appear in Table II. Qualitative interpretations are given in Table II.

Observed *T*_*m*_ values estimated from both Thermofluor and CD are significantly correlated with the same subset of mutational effects (Table I) which explain > 95% of the variation in *T*_*m*_ for both melting curves. The two melting steps share the most important predictor: the wild type F37 residue stabilizes both the unliganded open conformation against molten globule formation by ∼2.9 degrees, and the molten globule, relative to the denatured form by ∼2.2 degrees, almost irrespective of the variant in which it occurs (Figs. 8 (A),(B),(D). Other significant predictors involve I4, which stabilizes the native state vs the Molten Globule.

**Fig. 7.**
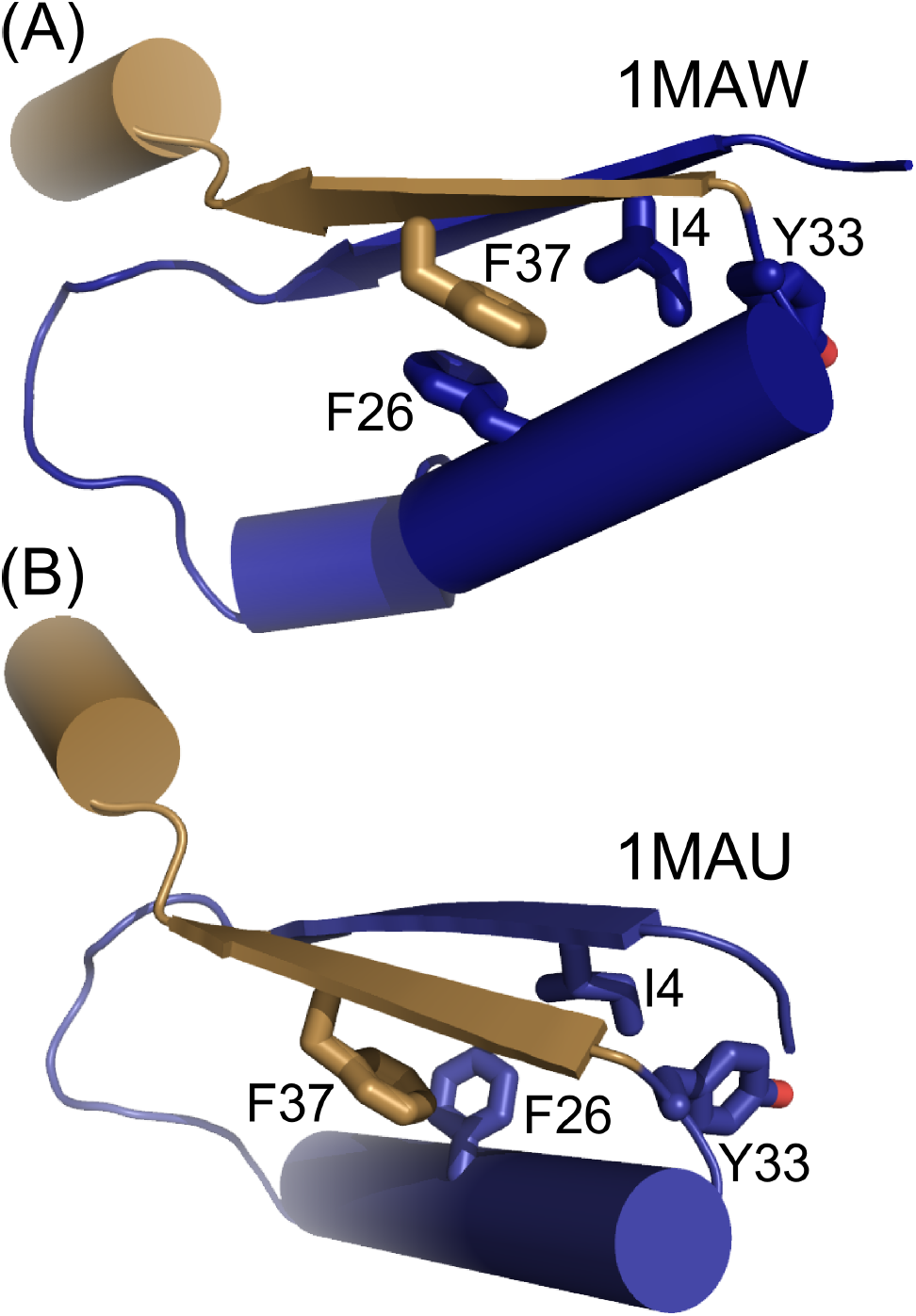
Wild type residue Phenylalanine 37 plays an inordinately large role in unliganded TrpRS stability. The significance of the large β_φ and β_θ coefficients (Fig. 6(C)) can be attributed, in part, to its interaction with the first crossover connection in the Rossmann fold. (A) In the unliganded state, F37 packs onto F26 in approximately a “T” configuration, holding the second β-strand (sand) against the surface of the first crossover connection (dark blue). (B) In the PreTS state, that interaction shifts to a less stable ^73^, “parallel-displaced” orientation.

**Fig. 8.**
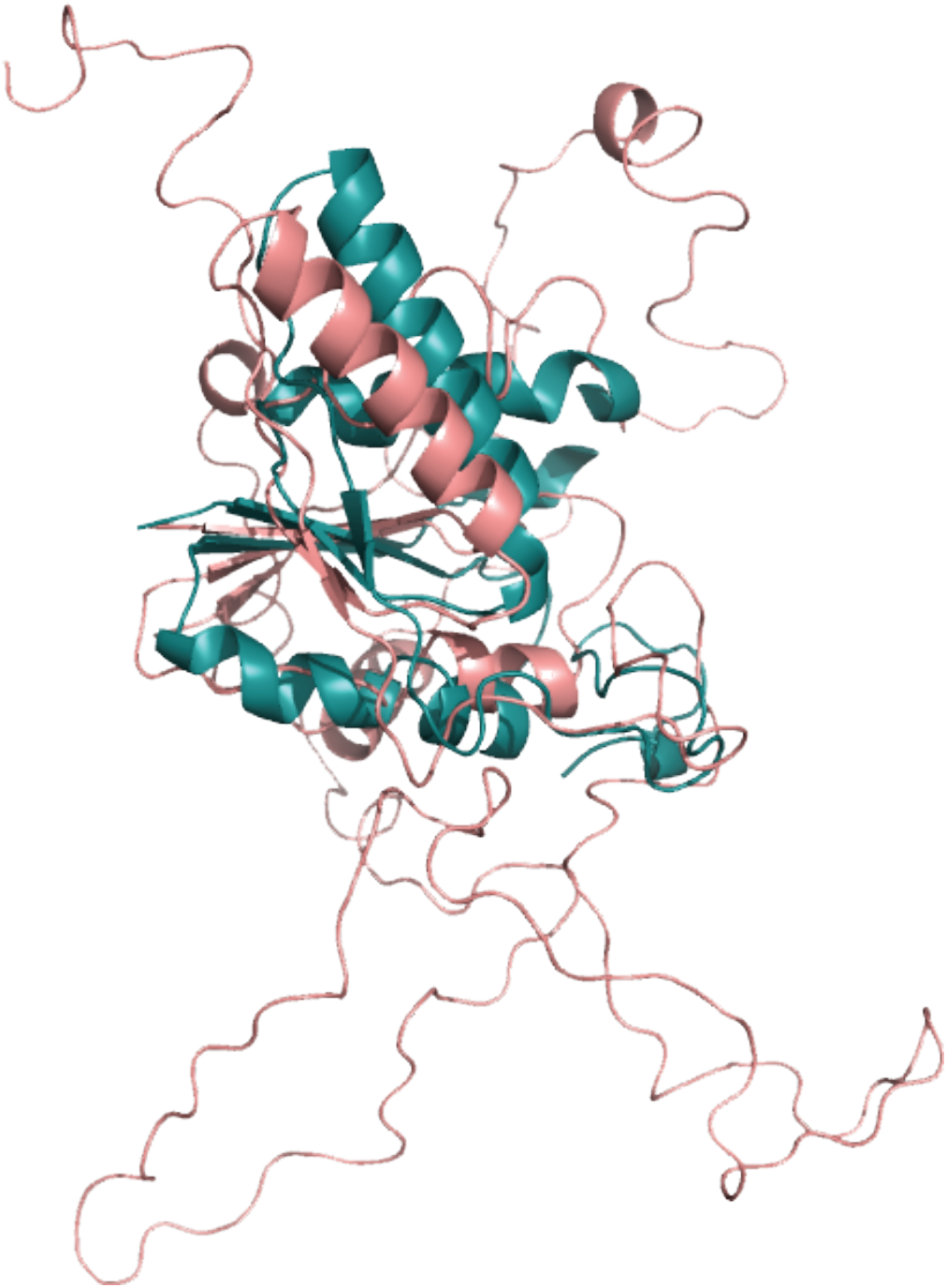
The TrpRS catalytic apparatus denatures last. The averaged structure from the ensemble at 94 C (salmon) is superimposed on the TrpRS Urzyme [teal ^24,25^]. The Urzyme remains largely intact whereas most of the monomer is essentially unfolded.

The F26*Y33 interaction has opposite effects, reducing 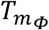 but increasing 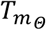. The net effect of the two residues (the main effects of F26 and Y33 plus their interaction), however, is to destabilize both the native protein and the molten globule by ∼ 0.5 degrees (see Fig. 1 inset). Student t-tests of these predictors are highly significant (Table I).

### Wild type residue F37 contributes inordinately to stability of both native and molten- globular configurations

Three of the mutational predictors have nearly the same effect on the stability of the native state and molten globule (Fig. 7(C)). Wild type residue F37 and the three-way interaction between the three aromatic wild type side chains exert both the native state relative to the molten globule, and on the molten globule relative to the fully denatured state. The two-way interaction between I4 and F37, the two side chains opposite one another on the two β strands is modestly destabilizing for the two states.

### The TrpRS Urzyme appears to be the last structure to denature

Haynie and Freire ^58^ outlined the structural energetics of molten globules in theoretical terms, refuting the widespread notion that there should be little or no enthalpy change on going from a molten globule to a fully denatured state. Steepness in the TrpRS CD transition implies that all or most of the α-helices melt cooperatively, with a large positive ΔH suggesting, in turn, that native-like tertiary packing also breaks down as the α-helices melt. Seelig and Schönfeld ^59^ discuss how Zimm-Bragg theory can consistently model enthalpy changes from multistate denaturation processes for which different probes suggest different melting temperatures. Discontinuities in the computational R_*g*_ and helical content profile coincident with heat capacity change maxima (Fig. 4) suggest that TrpRS unfolding may be amenable to similar treatment.

NMR measurements of residual dipolar coupling, RDCs, on oriented samples of denatured proteins ^60^ have been interpreted to mean that limited native-like structures are transiently retained in the denatured states of many proteins, suggesting that a continuum of such states leads from unfolded to native state. On the other hand, theoretical simulations ^61^ showed that transient native-like structure is unnecessary to account for non-vanishing RDCs. Correlation of both TrpRS melting processes to the same specific mutational effects in the D1 switch (Fig. 6) favors persistent native-like structures in the denatured state.

Supplementary Table SII provides both quantitative and qualitative insight into what that structure might be. All mutations show significant intrinsic and synergistic impact on both thermal transitions. F37 acts both independently and synergistically with F26 and Y33 to stabilize both native and molten globule states, and anti-synergistically with I4 and Y33 to destabilize both states. Several WT tertiary interactions—the intrinsic effect of wild type F37 and the Y33*F37 interaction—stabilize both native and molten globule states, elevating the temperature at which α-helices melt. The effects of F37 are consistent with the fact that this residue binds the second β-strand of the Rossmann fold to the first α-helix. That interaction is an important tertiary effect, as the N-terminus of that helix forms a rigid body with the anticodon-binding domain, relative to the Rossmann fold. Such stabilization probably plays a role in ensuring that none of the weaker helices melt until these tertiary interactions are broken, at which time most helices melt. In contrast, local interactions, F26*Y33 and I4*F26*Y33, have the opposite effect, destabilizing the native state, but not the molten globule (Fig. 6(D)). Several helices in the core of the protein remain intact at 94 C in the computational profile (Supplementary Fig. S4), whereas solvent exposed helices near the surface melt earlier.

Hilvert’s work with chorismate mutase ^62,63^ identified a potentially widespread efficient coupling between folding and catalysis—i.e. transition-state affinity. In light of those findings, it is curious that even the rather weak potentials constraining the simulations are sufficient to ensure that the last part of the TrpRS monomer to melt in the REX/DMD free energy simulations closely approximates the TrpRS Urzyme, a construct studied in the context of the evolution of aminoacyl-tRNA synthetases (Fig. 8). Thus, the modules that remain intact at the highest temperatures are also the most ancient part of the protein which, when excerpted, retain a full range of catalytic activities ^24,25,64^. We note the curious consistency between the structural integrity of the TrpRS Urzyme at high temperature and the experimental observation that the Urzyme itself seems to function as a molten globule ^65^, whose only possible well-folded state is its complex with the transition state for amino acid activation.

## CONCLUSIONS

### The detailed characterization of the TrpRS escapement mechanism motivates a need for experimental data on the coordinated thermodynamic impacts of D1 switch mutations on differential conformational stability

Differential scanning fluorimetry (Thermofluor; ^14^) is an attractive, high-throughput measurement. We compare Thermofluor measurements here with far UV circular dichroism melting curves for all 16 TrpRS variants arising from combinatorial mutagenesis of D1 switch residues.

### TrpRS unfolds via a molten globular intermediate

Sypro Orange fluorescence and θ_221_ melting curves are well-separated, linked and sequential. As with other proteins for which reversible multi-state melting behavior has been assumed ^49,50^, the product of the first process is the reactant for the second.

### Two novel kinds of evidence imply that Thermofluor melting temperatures contain useful information related to protein conformational stability

(i) Heat capacity changes and structural metrics from a single molecule computational free energy surface are consistent with an intermediate molten globular state of TrpRS as inferred from the experimental melting curves. (ii) T_m,φ_ and T_m,θ_ calculated from linear transformation of the combinatorial mutagenesis design matrix agree closely with the experimental values, implying that the experimental values also depend linearly on the design matrix as expected from the relationship of Calvin, Hermans, and Scheraga ^56^. These together argue that the multi-domain TrpRS structure, dimer dissociation, and aggregation have limited impact on the experimental melting transition temperatures. Thermofluor and CD melting curves therefore appear to be experimentally accessible indicators of conformational stability differences relevant to understanding conformational coupling. Mutational perturbation also increases substantially the resolution with which one can assess protein stability, affording access to structural information about folding transitions ^67-69^ and molten globular states that cannot be otherwise characterized.

### Tertiary packing and aromatic stacking interactions of F37 stabilize both the native state relative to the molten globule and the molten globule relative to the denatured state (Fig. 6 (A),(B))

The proportionality between experimental melting temperatures and those computed from the design matrix in Fig. 5(A) also suggests that all variants melt with very similar enthalpy changes, and that the mutation-dependent stability changes result mostly from perturbations to the lowest frequency domain motions that dominate the conformational entropy and have the greatest effect on catalysis.

### Ding, Jha, and Dokholyan ^70^ suggested the persistence of a core 3D structure in unfolded forms of many proteins

Remarkably, for TrpRS, that core, Fig. 8, coincides closely with the Urzyme, a model for the ancestral enzyme that itself appears to be a catalytically active molten globule ^65^.

### Establishing Thermofluor as a means of attributing stability to the mutated D1 switch side chains in this work will facilitate comparable analyses of other conformations from the structural reaction profile ^15,19,20,71^

This can be achieved using ligands that stabilize each conformation. The PreTS conformation can be stabilized by ATP and tryptophanamide, the products conformation by tryptophanyl-5’sulfamoyl adenosine, and a transition state by adenosine tetraphosphate ^19^. Analytical comparison of the ensemble of melting profiles will enhance the value of the combinatorial mutagenesis by enabling correlations with experimental ^1,3,26^ and computational ^6,28^ kinetic measurements.

## Supporting information

Supplemental tables, figures, text, and Matlab codes

## ACKNOWLEDGMENTS

Supported by NIGMS R01-40906 and R01-78227 to CWCjr and by NIGMS grants R01-GM114015, R01-GM064803, and R01-GM123247 (N.V.D). We gratefully acknowledge discussions with J. Hermans, to whose memory this work is dedicated, B. Lentz, G. Pielak, and A. Lee. D. Shortle’s suggestions helped improve coherence of the discussion, and the assistance of Ashutosh Tripathy in the UNC Macinfac facility. We thank A. Cook for suggesting to represent the combinatorial mutagenesis design matrix as a barcode.

## AUTHOR CONTRIBUTIONS

GW, VW, and CWCjr designed the experiments; GW wrote the Matlab software for data analysis; SNC and JD performed the REX/DMD calculations; SNC, JD, and ND analyzed the structural simulations; GW, VW, SNC, and CWCjr wrote and all authors approved the manuscript. None of the authors is aware of any conflicts of interest.

