## Supplemental tables, figures, text, and Matlab codes for "Thermodynamic impacts of combinatorial mutagenesis on protein conformational stability: precise, high-throughput measurement by Thermofluor"

^1.^ Department of Biochemistry and Biophysics, University of North Carolina at Chapel Hill, Chapel Hill, NC 27599-7260; ^2.^ Albert Sherman Center, AS4-2071, Program in Bioinformatics and Integrative Biology Worcester, MA - 01605, USA; ^3.^ Cystic Fibrosis and Pulmonary Diseases Research and Treatment Center, University of North Carolina at Chapel Hill, North Carolina, USA 27599-7248; ^4^ Departments of Pharmacology and Biochemistry & Molecular Biology, Penn State College of Medicine, Pennsylvania 17033, USA

**§1. Aggregation and reversibility**.

A persistent problem for stability studies, especially with multi-domain and/or oligomeric proteins, is that the various denatured states aggregate, limiting reversibility. This problem is compounded by the fact that the TrpRS dimer conceivably could denature by multiple pathways in which different subunits and different domains act either independently or in unknown, linked combinations. We present here the controls that bear on this question. Our conclusion is that aggregation does afflict the heating and cooling cycle necessary to demonstrate complete reversibility, but that the controls suggest that at the midpoint temperature of the transition, the folding is inherently reversible and that the aggregation results from the relatively high protein concentrations necessary for the experimental measurements.

**Relaxation rate determination after a temperature jump (Fig. S1A,B)**

A sample of native TrpRS comparable to those used in thermofluor stability assays was heated from 25 to 70 C and fluorescence readings were taken at time intervals. These data were corrected for background to give Φ_t_. An asymptotic value for the total amplitude, Φ_0_, was assumed, and differences, (Φ_0_ – Φ_t_) were plotted on semi-log paper, adjusting the Φ_0_ value to eliminate curvature in the plot. Once a suitable value for Φ_0_ was determined, the slope, k, was estimated by least squares, with a statistical error of slightly more than 1%.

We estimated the rate constant for Thermofluor melting from a temperature jump experiment (Fig. S1A). Fluorescence readings fitted to a single exponential yielded a rate constant of 0.0033/s, which corresponds to a half-time of 209 seconds or 3.5 minutes for relaxation at 70^o^ C. The 21-minute time course for the melting curve for the interval 25-70 C is six half-lives, allowing ~98% of the molecules to equilibrate. Temperature is increased in 0.46 minute/degree steps.

To ensure that this rate does not affect the melting curve, we measured the melting curve using ANS as the dye in a spectrofluorimeter with steps allowing 1.5 minutes/degree. Curves for the two different dyes and time-courses (Fig. S1B) are essentially indistinguishable, with <$T_{m}$> = 69.8 ± 0.4 degrees. The measurements therefore allow sufficient equilibration time.

**Recovery of near UV CD (Fig. S1C)**

Efforts to reverse aggregation induced by Thermal unfolding of TrpRS α-helices had limited success. Molten globule formation, however, can be extensively reversed. We emulated a procedure (1) that cycles through a rapid increase to the melting temperature until a steady state is achieved, followed by cooling to 25^o^ C. Owing to artefactual fluorescence changes observed in Sypro Orange fluorescence in small plastic wells after rapid temperature jumps, we assayed this process using near UV CD in the absence of dye, which for TrpRS is very weak (Fig. 3A). Nonetheless, the Cotton effect at 270 nm is regenerated on cooling. The recovered Cotton effect is more pronounced than that of the starting material, suggesting that some starting material may be incorrectly folded and not contribute to the near UV CD. To validate this interpretation, we assayed the enzyme by active-site titration (2) before and after heating and cooling. After cooling 40% of the protein had precipitated; the 60% of soluble protein recovered ~80% of the original active-site titer (Fig. 3B). Thermofluor melting thus can be reversed.

The near UV CD spectrum for an ~80 μM wild-type TrpRS sample was determined (Fig. 3A). Then, the sample was heated from 25 C to 68 C for 10 minutes and another spectrum was recorded. The sample was then returned to 25 C and a third spectrum recorded. No attempt was made to clarify the solution by centrifugation between spectra, nor was any correction made for loss of aggregated material. Evidence of structural recovery is clear, but difficult to quantitate, owing to the weakness of the signal.

**Active-site titration (Fig. S1D)**

Active-site titration is a single-turnover experiment that is the gold standard for measuring the fraction of active enzyme molecules (2, 3). Slightly less than stoichiometric amounts of enzyme are incubated with radiolabeled γ-^32^P ATP and tryptophan and the loss of label is measured at time intervals spanning the first-order reaction and subsequent turnover. This experiment is useful if and only if there is a pre-steady-state burst. Aminoacyl-tRNA synthetases retain the activated amino acid intermediate, and therefore exhibit burst kinetics. That burst is proportional to the fraction of active sites contributing to the reaction.

**Perturbation of the molten globular state by dye binding (Fig. S1E)**

It is important to distinguish two possible dye-binding modes, opportunistic and/or causative. Does the dye simply detect molten globule formation, or does it bind tightly enough to influence the temperature at which it forms? The weak TrpRS Cotton effect at 270 nm makes it technically difficult to follow molten globule formation with and without dye in this case. The model in the center of Fig. 5 provides the basis for an indirect test of such effects. To help assure that comparisons between Thermofluor and θ_221_ are unaffected by the presence of dye in the former experiments, we compared the θ_221_ melting behavior with and without Sypro Orange at the concentration used in Thermofluor experiments (Fig. S2E). Fitted $T_{m}$s (78.15 ± 0.1^o^C; 77.93 ± 0.1^o^C for WT TrpRS plus and minus Sypro Orange) are within experimental error.

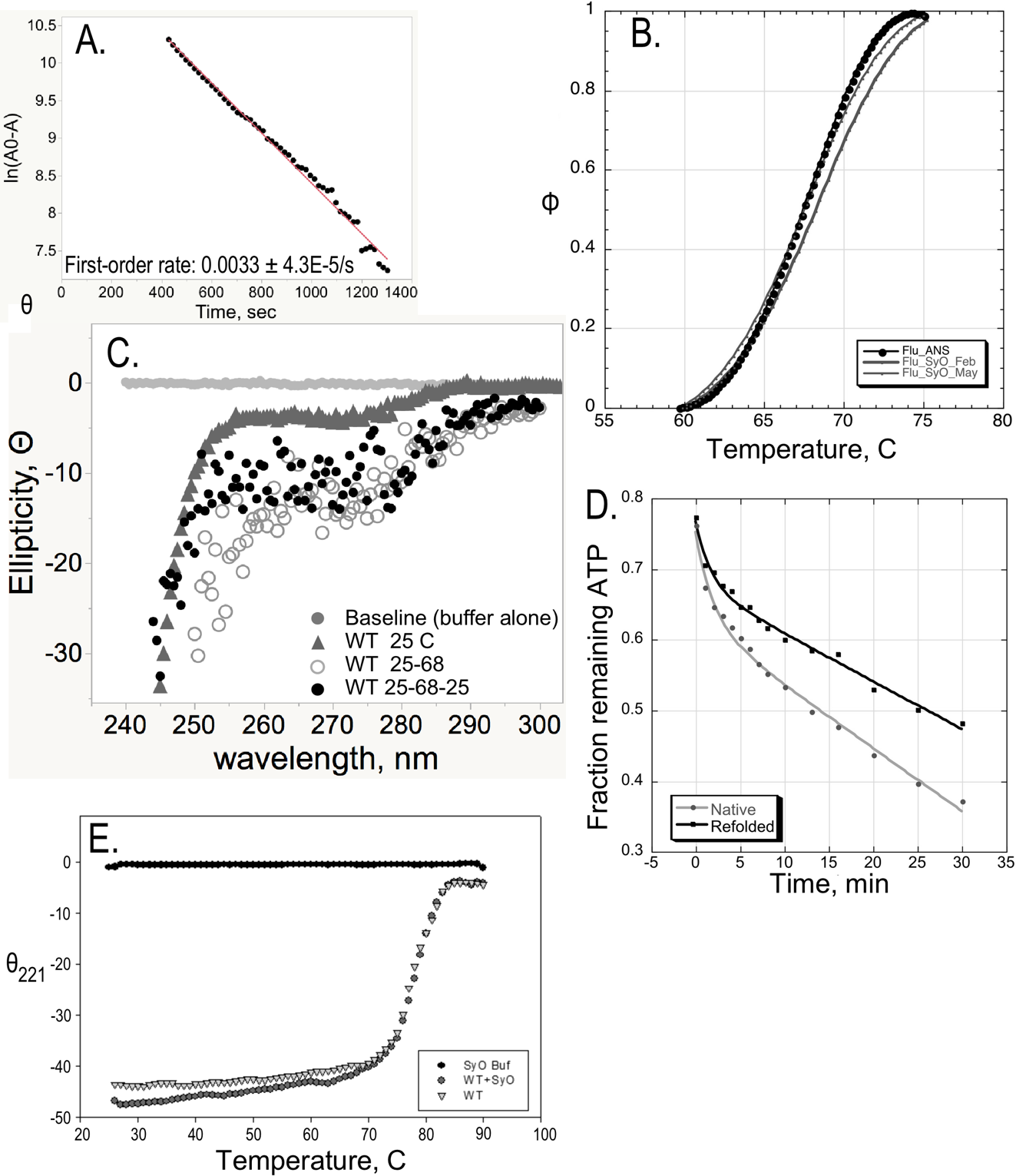

Fig. S2. Control Experiments. Details are described in §1. A. Estimation of the first-order relaxation rate at 70 C after a temperature jump from 25 C. B. Normalized melting curves for ANS (Black circles) and Sypro Orange (grey) performed with different equilibration times. Fitted parameters for the three curves are indistinguishable. C. Recovery of the Cotton Effect at 270 nm for an 80 μM TrpRS solution on cooling after a temperature jump to 70 C. D. Active-site titration of native and recovered TrpRS for the experiment in C. E. Sypro Orange does not significantly perturb the melting behavior detected for native TrpRS by θ_221_.

**§2. Protein concentration effects.**

Thermofluor (6 μM) and CD (8 μM) measurements were done at slightly different protein concentrations because the signal to noise was optimal at the respective concentrations. Thermofluor melting experiments, cannot be done far away from the concentration we chose, without serious reduction in signal to noise. As a control, we measured the CD melting curve at a concentration of 2 μM. The melting curves are nearly indistinguishable (Fig. S2). Four-fold dilution decreased T_m_ by about a degree, which is more than 6 standard errors and probably significant. However, the only volume changes other than pipetting errors in these experiments were those caused by thermal expansion of the buffer as temperature increased, which were both minimal and the same for all samples. The protein concentration was 10^3^ times K_D_, so the extent of dimer dissociation due to dilution is therefore expected to be negligible, because volume changes are negligible. Moreover, differences between Thermofluor and CD melting arising from the slight protein concentration difference would be expected to be the same for all mutants, leaving our conclusions unaffected.

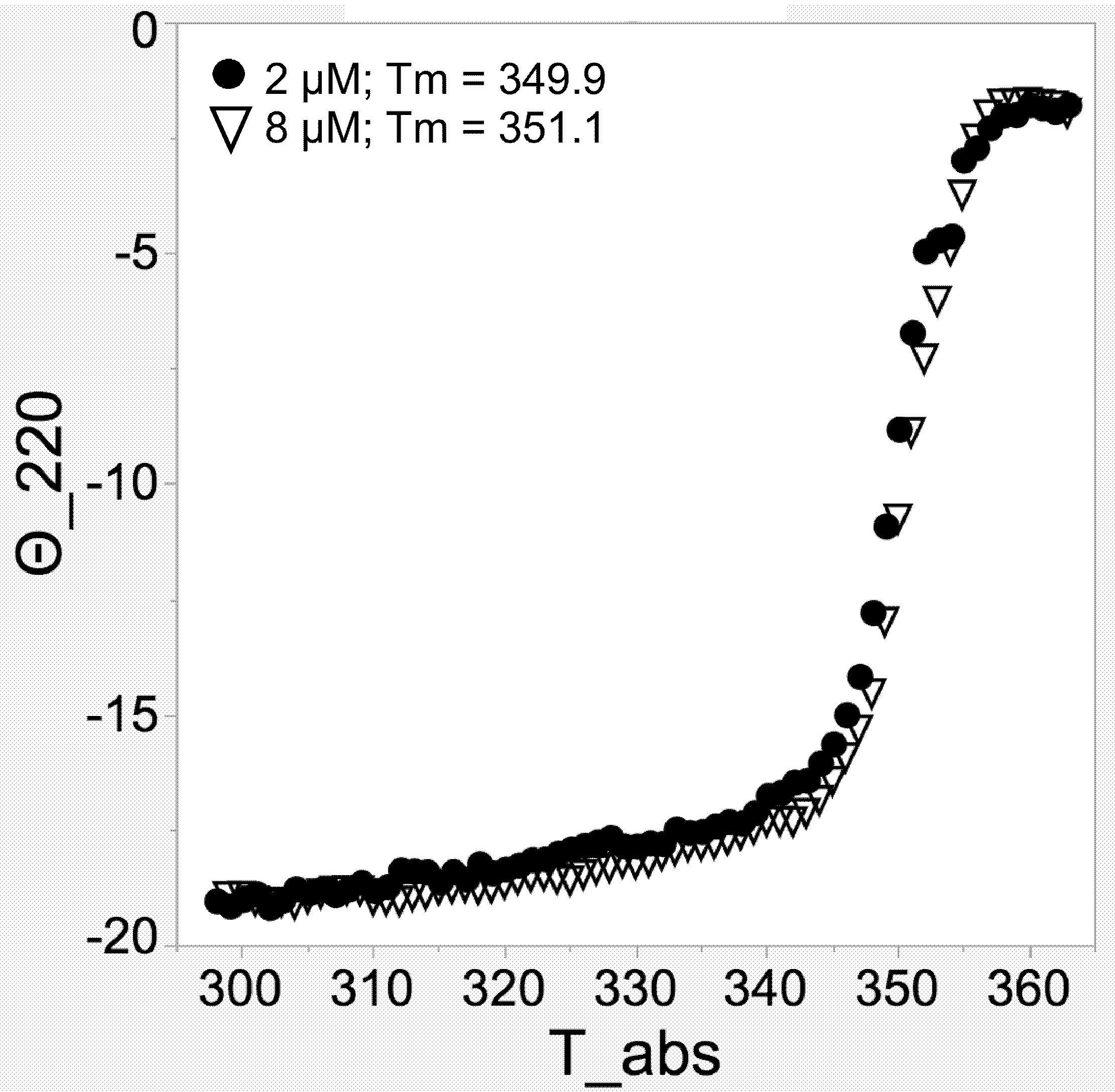

Fig. S1. Protein concentration dependence of thermal melting. CD melting curves at 221 nm of native TrpRS at the concentration used for all experiments (8 μM, inverted open triangles) and at a four-fold dilution ((2 μM; black circles). T_m_ values estimated by the ratio method are included in the key.

**§3. Melting Curves**.

Much of the manuscript is devoted to validating thermodynamic interpretations of unfolding studies of the 16 combinatorial variants of *B. stearothermophilus* tryptophanyl-tRNA synthetase. Specifically, we identify two unfolding transitions—molten globule formation and loss of α-helical secondary structure—and show that the combinatorial mutants induce correlated effects on the stabilities of the three states related by the two transitions. Experimental profiles of the two transitions are compared in Fig. S2A and B, from which it can be seen that the transitions from one state to another can be represented to good approximation by linear temperature dependent profiles outside the transition region linked by a Boltzman type sigmoidal curve representing the transition between them Fig. S3.

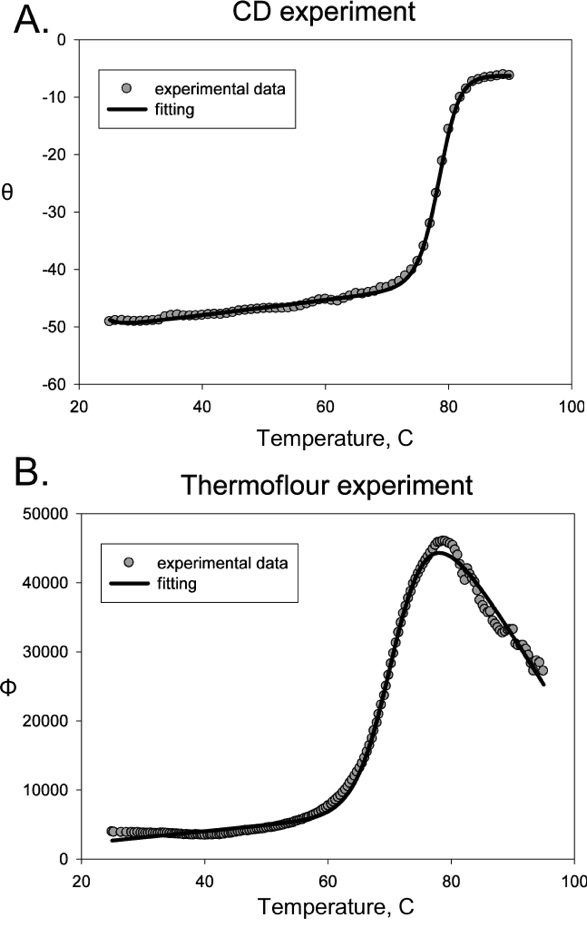

Fig. S2. *Melting curve analysis*. Fitting of representative CD (A) and Thermofluor (B) melting curves to the thermodynamic two-state model (eqs. 3, 4). C. Decomposition of the experimental signal within the transition into fractional probabilities of States n and n+1 by the ratio method.

**§4. Estimating temperature-dependence of successive state probabilities.**

*The Ratio Method (Fig. S2)*. We based the “ratio” method on the experimentally observed fact that the initial (P_1_=1 and P_2_=0) and final (P_1_=0 and P_2_=1) parts of denaturation curves are linear, where P_1_ and P_2_ are the probabilities of the two states. This method also assumes a reversible equilibrium between two (quasi-) steady states (P_1_ and P_2_). As we have been unable to show cleanly that TrpRS melting behavior is fully reversible, we justify this assumption by showing that computational free energy simulations reproduce the volume, heat capacity and helix content changes implied by the two observed thermal transitions.

Consider a ratio of differences between fluorescence changes for states S1 and S2 and extrapolations of final and initial slopes, shown graphically in Fig. S2:

$$F_{A}=F_{ini}=a_{1}+b_{1}T$$

$$F_{B}=F\left( T \right)=\left( a_{1}+b_{1}T \right)P_{1}+\left( a_{2}+b_{2}T \right)P_{2}$$

$$F_{C}=F_{end}=a_{2}+b_{2}T$$

(S2)

F_B_, the fluorescence at a given temperature, is thus the sum of the fluorescence from State 1 (F_B_ - F_C_) and from State 2 (F_A_ - F_B_). Their ratio is therefore an apparent equilibrium constant for reaction at that temperature:

| .$F_{A}-F_{B}=\left( \left( a_{1}+b_{1}T \right)-\left( a_{2}+b_{2}T \right) \right)P_{2}$ | (S3a) |
| --- | --- |

$$F_{B}-F_{C}=\left( \left( a_{1}+b_{1}T \right)-\left( a_{2}+b_{2}T \right) \right)P_{1}$$

(S3b)

$$\frac{F_{A}-F_{B}}{F_{B}-F_{C}}=\frac{P_{2}}{P_{1}}=e^{-\frac{\Delta G\left( T \right)}{RT}}$$

(S4)

since $P_{1}=\frac{1}{1+e^{\frac{-\Delta G}{\mathrm{RT}}}}$ and $P_{2}=1-P_{1}$

Equation (4) leads to a direct determination by interpolation of $T_{m}$, the melting temperature where the two, apparent, state probabilities are equal and ΔG*_app_* = 0. Note, that Eq. 4 was derived without any model for the Gibbs energy ΔG*_app_* and relies only on the linearity of the initial and final parts of the melting curve and the quality of the sigmoidal fit.

Fitting the two melting processes affords quantitative metrics supporting a three-state model for TrpRS thermal unfolding (Fig. 5; Fig. S2). We assume that TrpRS denaturation proceeds through three states, whose probabilities total unity:

$$P_{1}+P_{2}+P_{3}=1$$

(S5)

where

$$P_{1}=\frac{e^{-\frac{G_{1}}{kT}}}{e^{-\frac{G_{1}}{kT}}+e^{-\frac{G_{2}}{kT}}+e^{-\frac{G_{3}}{kT}}};P_{2}=\frac{e^{-\frac{G_{2}}{kT}}}{e^{-\frac{G_{1}}{kT}}+e^{-\frac{G_{2}}{kT}}+e^{-\frac{G_{3}}{kT}}};$$

$$P_{3}=\frac{e^{-G_{3}/kT}}{e^{-G_{1}/kT}+e^{-G_{2}/kT}+e^{-G_{3}/kT}}$$

(S6)

or

$$P_{1}\left( {T_{m}}_{\Phi} \right)=P_{2}\Rightarrow\Delta G_{fluo}=0$$

(S7)

Transitions connect two states in Thermofluor and again in the CD melting curves. Normalized Thermofluor and CD melting curves for the same variant afford direct estimates of the three state probabilities as functions of temperature (Fig. 6). As experimental free energy differences reflect state probabilities, ($\Delta G_{fluo}=G_{2}-G_{1}$) and ($\Delta G_{CD}=G_{3}-G_{2}$), the experimentally determined values lead to revised expressions for probabilities, P_i_ (see Eq.(6)):

$$P_{1}=\frac{1}{1+e^{-G_{fluo}/kT}+e^{-\left( \left( G_{3}-G_{2} \right)+\left( G_{2}-G_{1} \right) \right)/kT}}=\frac{1}{1+e^{-G_{fluo}/kT}+e^{-(G_{CD}+G_{fluo})/kT}};$$

$$P_{2}=\frac{1}{1+e^{{-G}_{fluo}/kT}+e^{-G_{CD}/kT}};$$

$$P_{3}=\frac{1}{1+e^{-\left( \left( G_{1}-G_{2} \right)+\left( G_{2}-G_{3} \right) \right)/kT}+e^{-\left( G_{2}-G_{3} \right)/kT}}=\frac{1}{1+e^{(G_{CD}+G_{fluo})/kT}+e^{G_{CD}/kT}}$$

(S8)

State probabilities based on ΔG_fluo_ and ΔG_CD_ are then:

$$P_{1}=\frac{1}{1+e^{-{\Delta G}_{fluo}/kT}+e^{-({\Delta G}_{CD}+\Delta G_{fluo})/kT}};$$

$$P_{2}=\frac{1}{1+e^{\Delta G_{fluo}/kT}+e^{-{\Delta G}_{CD}/kT}};$$

$$P_{3}=\frac{1}{1+e^{(\Delta G_{CD}+\Delta G_{fluo})/kT}+e^{{\Delta G}_{CD}/kT}}$$

(S9)

Intersections $P_{1}=P_{2}$ and $P_{2}=P_{3}$ explicitly determine ${T_{m}}_{\Phi}$ ($P_{1}\left( {T_{m}}_{\Phi} \right)=P_{2}\Rightarrow\Delta G_{\Phi}=0$) and ${T_{m}}_{\theta}$ ($P_{2}\left( {T_{m}}_{\Theta} \right)=P_{3}\Rightarrow\Delta G_{\theta}=0$), respectively, using Eqs. (8). This is not true for the temperatures at which P_2_ reaches a maximum and where P_1_=P_3_. Thus, the maximum value of $P_{1}$ and values of $P_{1}$($P_{2}$=max) = $P_{3}$($P_{2}$=max) and discrepancies between these two temperatures are quantitative metrics of the separation of the two linked processes in TrpRS denaturation. For WT and all 15 mutant TrpRS, these metrics closely approach 1.0 (0.95±0.02); 0.0 (0.025±0.011); and 0.0 (0.18±0.09); (e.g. Fig. 5). This analysis therefore confirms that the population of denatured α-helices is nil until the population of native molecules is essentially exhausted and that of molten globules reaches a maximum.

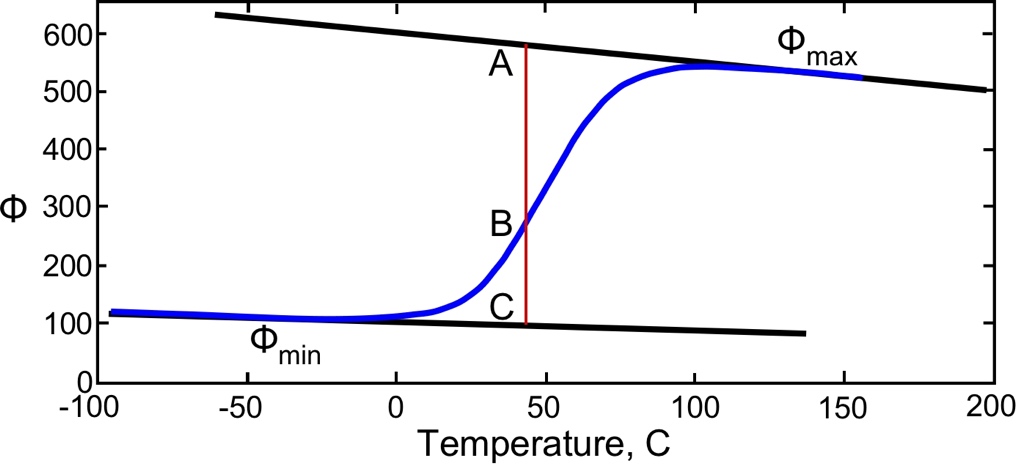

Fig. S3. *Decomposition of the experimental signal within the transition into fractional probabilities of States n and n+1 by the ratio method*.

**§5.** **Regression modeling**

The 2^4^ factorial design (4) of sixteen variants balances comparisons of mutant behaviors. The intrinsic effect of each mutation is averaged over eight contexts, that of each double mutant over four contexts, and each triple mutant over two different contexts (wild-type and mutant in the remaining residue). Regression modeling uses this averaging to enhance the precision and sensitivity of mutational analysis, quantitation, and significance testing of high-order interactions (see supplement to ref 5)).

Misconceptions nonetheless persist arising from confusion between the actual experimental observations for individual multiple mutants themselves, and the coupling energies they imply. Energetic coupling between residues depends on how the effect of a particular mutation changes in different contexts (wild-type, mutations at other sites, etc.) in which it is observed. For that reason, the entire set of combinatorial mutants must be analyzed as an ensemble. Comparison between a quadruple mutant and WT protein may suggest only a small interaction when, in fact, the overall coupling can be quite significant. Our own work (5, 6) and that of others (7) provide examples of how higher-order coupling emerges from multi-mutant thermodynamic cycles.

**§6.** **Thermodynamic cycle analysis of combinatorial mutants**

The longer-term goal of this project is to analyze how these induced stability changes are altered in different conformations along the structural reaction profile, which can be induced by particular combinations of ligands. To that end, it is useful to construct probability distributions for the three distinct states, as outlined in the section “**TrpRS denatures via an intermediate, molten globular state**” and illustrated in Fig. S2 and Fig. 6 of the paper.

State probability distributions for the 16 variants are compiled for reference in Fig. S4.

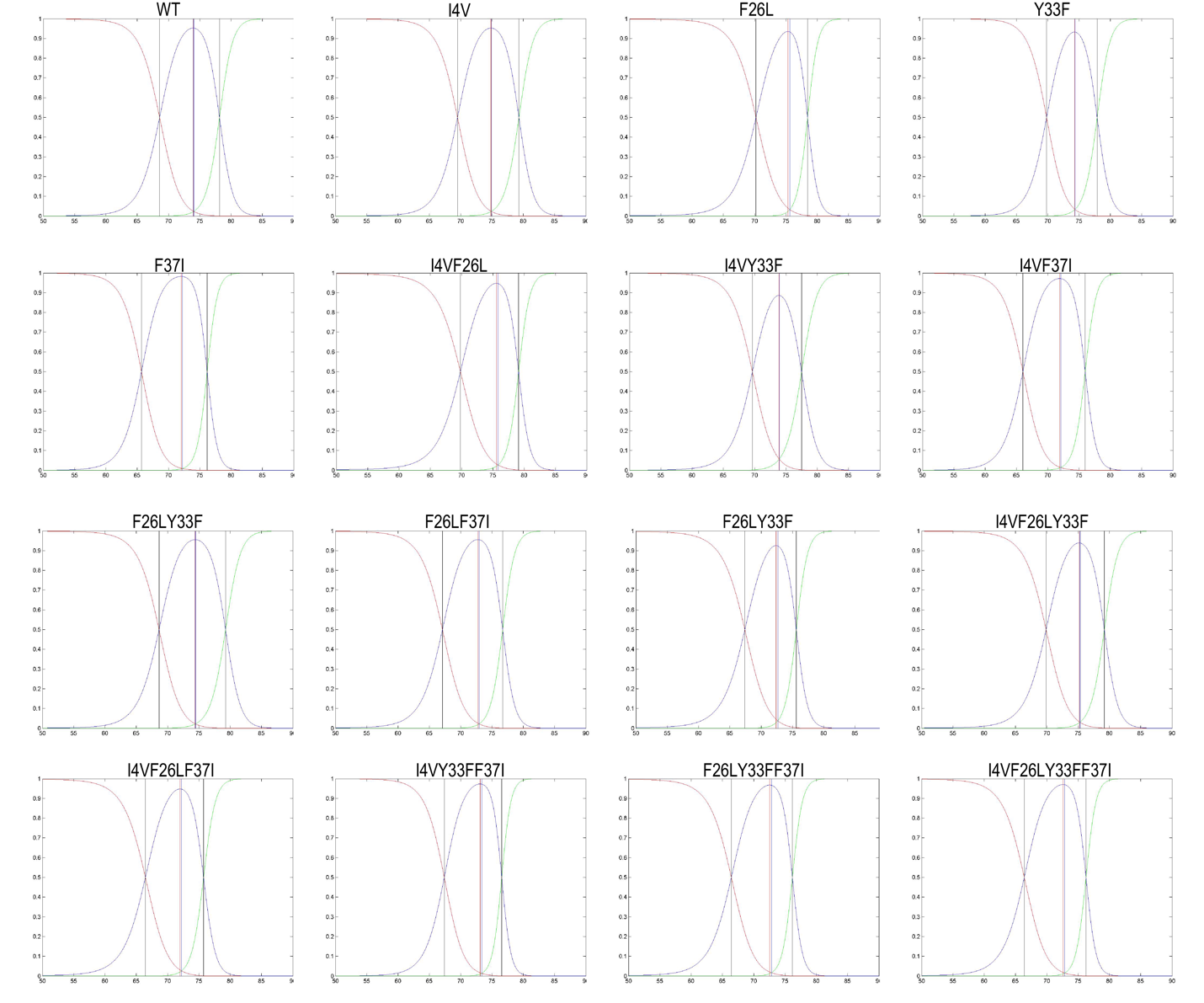

Fig. S4. *State probability distributions constructed on the basis of Thermofluor and CD unfolding curves, as described in the text of the paper*. The variants are ordered sequentially from Wild Type to the quadruple mutant, each variant ordered according to its position in the sequence. Black vertical lines denote temperatures at which the probabilities of two successive states are equal, and for which the free energy of the transition is therefore 0; red vertical lines denote the temperatures at which the molten globule probabilities are maximal; and blue vertical lines denote temperatures at which the native and fully unfolded states have equal probabilities. Note that the blue lines have uniformly low probabilities, highlighting that the succession of states in thermal unfolding is nearly disjoint, and that the differences in temperature between the maximum molten globule probability and that for which the native and unfolded states are in equilibrium nearly coincide.

**§S7. The effect of divalent metals**

Divalent metals somewhat surprisingly destabilize the native TrpRS conformation by varying amounts relative to the molten globular state. Thermofluor melting temperatures are summarized in Fig. S5(A) for four conditions: 20 mM MgCl_2_, the condition that was fully replicated for TrpRS preparations performed on two occasions separated by three months, 1 mM MgCl_2_, 1 mM MnCl_2_. As no divalent metals were added to the CD melting curves, we performed melting curves for all variants without any added metals, to obtain the plot shown in Fig. S5(B). It should be noted that the multiple regression equations used to fit the data in Fig. S5(A) included only one parameter for each metal condition, that parameter was equivalent to an offset along the y axis. Because the R^2^ value is so high (R^2^ = 0.94, it is unlikely that the metals exert distinguishable effects on different combinations of mutations.

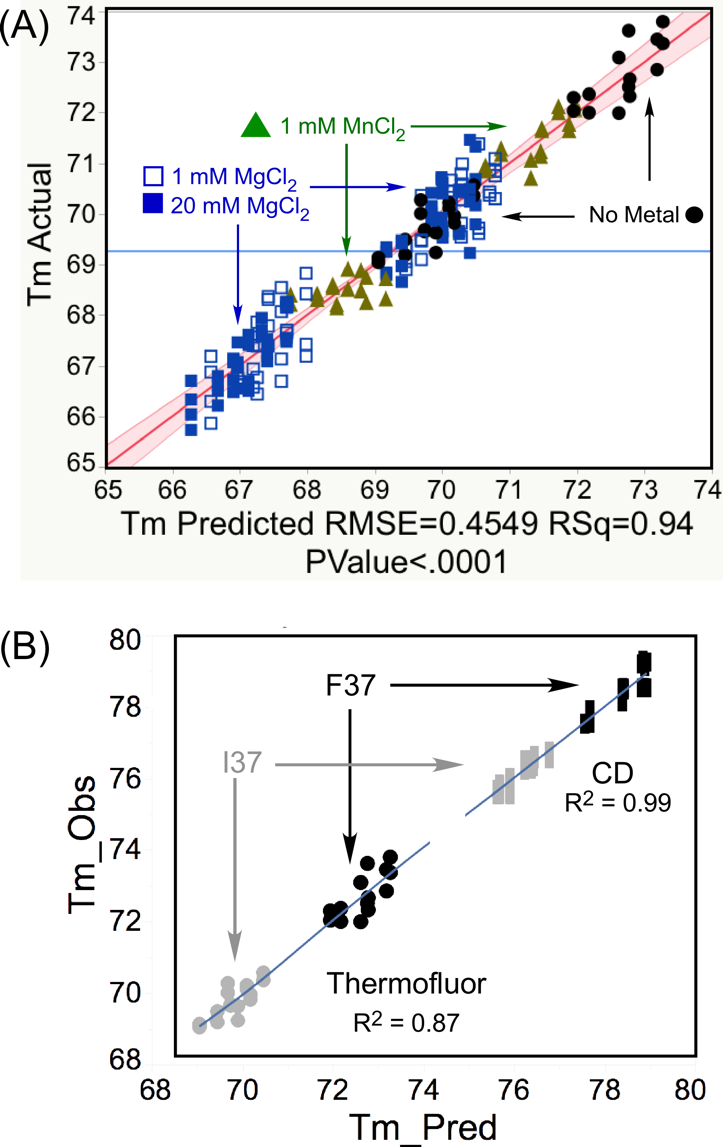

Fig. S5. Thermofluor melting temperatures at different metal concentrations. (A) All Thermofluor measurements, fitted to a single multiple regression model with one variable to denote each separate metal configuration. (B) Metal-free melting temperatures plotted together with CD melting temperatures. Compare with Fig. 5(B) of the main text.

**§S8. Physico-chemical helix propensities**

In order to assess the relative tendency of sequences along the TrpRS primary structure, the propensity for helix formation was determined as a function of temperature using the AGADIR server (8). These are plotted in Fig. S4, where they are also shaded according to the domain (Rossmann fold or anticodon-binding domain) to which they belong. A threshold value of 2.5 separates the helices with the strongest helical propensities, showing that both domains contain sequences with exceptionally high propensity.

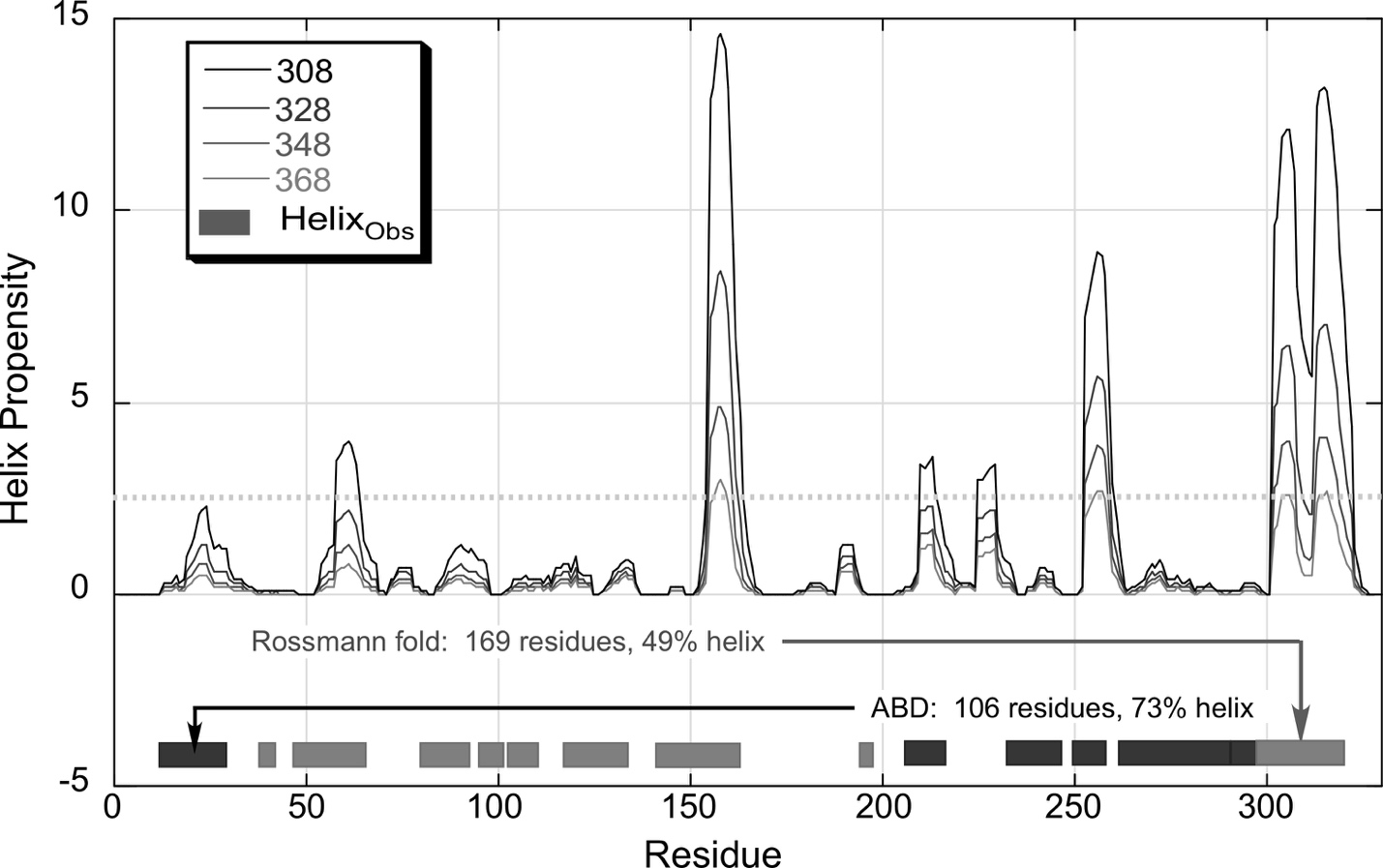

Fig. S6. *TrpRS helix propensities computed using the Agadir server (34) for different temperatures*. Only three of the helices observed in the TrpRS structure (residues 145-165; residues 233-246; and residues 303-318) have peak propensities exceeding 2.5 (dashed line) at the highest temperature used in our experiments (368 K). The Rossmann fold (gray) and anticodon-binding (ABD; black) domains are interleaved, as the first helix participates in both, as part of the Rossmann fold that actually moves as a rigid body with the ABD. This interleaving likely contributes to the high cooperativity of the helix melting transition monitored by θ_221_.

The dependent variables for the experimental design matrix for thermodynamic cycle analysis are given in Table S1, in which bold-face entries indicate minimum and maximum values for both melting temperatures. This table also indicates the highly reproducible nature of the measurements.

Significant main effects and higher order coupling interactions plotted as histograms in Fig. 6 of the paper are tabulated in Table SII, together with their Student t-test P-values.

**§9 Supplementary Tables**

Table SI. $T_{m}$ values obtained by Thermofluor and CD, estimated by model fitting and model-free analysis. Each combinatorial variant has a different row. Values derived from replicated Thermofluor melting curves are averages, accompanied by their Standard Deviations, calculated by the Excel Stdev function. The minimum and maximum Tm values for the two successive transitions are in bold face to emphasize the respective ranges.

| Variant | Tm_flu, Model | Stdev_Model | Tm_flu, Ratio | StdevRatio | Tm_CDModel | Stdev_CDModel | Tm_CD_ratio | Stdev_CD_ratio |
| --- | --- | --- | --- | --- | --- | --- | --- | --- |
| WT | 69.06 | 0.36 | 69.19 | 0.23 | 77.94 | 0.28 | 77.90 | 0.20 |
| I4V | 69.97 | 0.16 | 70.14 | 0.15 | 78.79 | 0.72 | 78.36 | 0.73 |
| Y33F | 70.45 | 0.21 | 70.28 | 0.23 | 78.15 | 0.46 | 78.09 | 0.69 |
| F26L | **70.63** | 0.30 | **70.29** | 0.18 | 77.53 | 0.55 | 77.28 | 0.79 |
| F37I | 66.03 | 0.34 | 66.32 | 0.35 | 75.84 | 0.54 | 75.91 | 0.54 |
| I4V_F26L | 70.36 | 0.69 | 69.86 | 0.69 | **78.83** | 0.41 | **78.48** | 0.26 |
| I4V_Y33F | 70.53 | 0.41 | 70.09 | 0.36 | 77.25 | 0.38 | 77.36 | 0.38 |
| I4V_F37I | **66.41** | 0.17 | **66.56** | 0.16 | 75.74 | 0.41 | 75.68 | 0.36 |
| F26L_Y33F | 69.07 | 0.38 | 69.05 | 0.34 | 78.65 | 0.85 | 77.95 | 0.48 |
| F26L_F37I | 67.25 | 0.10 | 67.38 | 0.15 | 76.51 | 0.36 | 76.33 | 0.16 |
| Y33F_F37I | 67.87 | 0.16 | 67.80 | 0.15 | 75.69 | 0.00 | 75.60 | 0.09 |
| I4V-F26L-Y33F | 70.21 | 0.23 | 69.99 | 0.18 | 78.40 | 1.13 | 78.05 | 0.85 |
| I4V-F26L-F37I | 67.58 | 0.18 | 67.81 | 0.20 | 76.58 | 0.10 | 76.47 | 0.14 |
| I4V-Y33F-F37I | 66.95 | 0.30 | 67.11 | 0.28 | **75.44** | 0.58 | **75.27** | 0.54 |
| F26L-Y33F-F37I | 66.77 | 0.23 | 66.99 | 0.20 | 75.94 | 0.43 | 75.86 | 0.29 |
| I4V-F26L-Y33F-F37I | 66.81 | 0.17 | 67.21 | 0.17 | 76.28 | 0.06 | 76.27 | 0.05 |

Table SII. Interpretation of mutational effects on relative stability of native and molten globular TrpRS (See also Figure 6A). Bold face denotes more substantial effects.

| **Predictor** | **Native** | **Molten globule** | **Notes** |
| --- | --- | --- | --- |
| **(WT) F26** | Little effect | **Destabilizes** |  |
| **(WT) Y33** | Stabilizes | No Effect |  |
| **(WT) F37** | **Stabilizes** | **Stabilizes** | pi-stacking across secondary structures changes from one conformation to another. |
| **(WT) I4*F26** | **Stabilizes** | No Effect | Side chains interact across secondary structures. |
| **(WT) F26*Y33** | **Destabilizes** | Little effect |  |
| **(WT) F26*F37** | Little Effect | Destabilizes |  |
| **(WT) Y33*F37** | Destabilizes | No effect |  |
| **(WT) I4*F26*Y33** | **Destabilizes** | No effect |  |
| **(WT) I4*Y33*F37** | **Stabilizes** | Destabilizes |  |
| **(WT) F26*Y33*F37** | **Stabilizes** | **Stabilizes** |  |
| **(WT) I4*F26*Y33*F37** | **Destabilizes** | No effect | This effect accelerates both amino acid activation (5) and acylation (9). |

**§10. MATLAB codes**

**Part 1. Data entry**

```
% Part 1: reading the melting data file in original format and converting it into matrix file
% Input: original melting data (OriginalData.txt).
% Output: text file (data.txt) containing matrix with following colums:
% well number, ligand concentration, temperature and fluorescence reading

clear all
tic
disp('-----------Start--------------------------')

%--------------Parameters to enter manually------------
%ligand concentrations
% if concentration changes along rows and repeats along columns then O=1
% if concentration changes columns rows and repeats along along then O=2
O=2;

if O==1 % concentrations change horizontally along with well numbers
    nCl=6;
    Nr=7;
elseif O==2 % concentrations change vertically and repeats horizontally
    nCl=7;
    Nr=6;
end

Ncol=9; % number of columns in the original data file.
%Might be different for different experiments

%-----------Ligand concentrations----------------------------
Cl=[250;600;1400;3500;8200;13000;20000];%concentrations can be defined in this line too or below via dialog (in uM)
Cl=Cl*10^(-6);%turn into Molar concentration

Ct=input('Do you want to input ligand concentrations (in M) (y/n)? ','s');

if Ct=='y' || Ct=='Y'
    for i=1:nCl
        Concentration=['Concentration(',num2str(i),')='];
        Clt = input(Concentration, 's');
        Cl(i)=str2double(Clt);
    end
end

%saving the ligand concentrations in 'Concentrations.txt' file for further
%use in refits
save LigandConcentrations.txt Cl -ascii


[Conc,cl]=size(Cl);% Conc= number of ligand concentrations

% error check
if Conc~=nCl
    error('Wrong set of concentrations')
end

%% --------------------Part 1-------------------------------
% Reading the original data and saving only the data in a new file
fid = fopen('OriginalData.txt');
file = textscan(fid, '%s','delimiter', '\n');

%textscan returns a 1-by-1 cell array,file that contains a 31-by-1 cell
%textscan reads the file as a string, line-by-line
lines=file{1};
L=length(lines);% total number of lines in the original data file
fclose(fid);


fid = fopen('OriginalData.txt');

A=zeros(L,Ncol);% prelocate memory for data file
i=0;% index of the row in the new data file

kk=0;% index for the data row in the well

nn=0;% index for repeats
nc=1;% index for the value of concentration
s=2;% index for lines: s=2 "Empty line", s=1 line to save, s=0 start counting measurements in the new well

%checking line-by-line
for l=1:L% index for line in the file

    tline = fgetl(fid);
    B=str2num(tline);

    firstword = sscanf(tline,'%9c', 1);

    %Start saving data from the next line
    if ischar(firstword) && strcmp(firstword,'Well	Time') && s==2
       s=0; % start counting measurements in the new well
       kk=kk+1; % number of wells for analyses

    elseif s==0 %&& i>0

        if ischar(tline) && length(B)==Ncol %&& i>0 % saving data
           i=i+1;% index of the row in the new data file
           A(i,:)=B; %Adding next data row, B

        elseif isempty(B) || B(1)~=A(i-1,1)% Finding the end of the data in the well
            s=2;
        end
    end


end
fclose(fid);

% deleting empty (zero-containing) rows
for k=i+1:L
    A(i+1,:)=[];
end


I=i; % number of lines in the data file
DD(:,[1,3])=A(:,[1,3]);% well and temperature
DD(:,2)=0;% ligand concentration
DD(:,4)=A(:,9);% fluorescence readings


%% --------------------Part 2-------------------------------
% Reading the original data and saving only the data in a new file
if O==1   % concentrations change horizontally along with well numbers
  nc=1;
  DD(1,2)=Cl(1);

  for ik=1:I-1
      if DD(ik+1,1)==DD(ik,1)
          DD(ik+1,2)=Cl(nc);
      else
          nc=nc+1;

          if nc>Conc
              nc=1;
              DD(ik+1,2)=Cl(nc);
              %error('something wrong with concentrations arrangements')
          else
          DD(ik+1,2)=Cl(nc);
          end
      end
  end

elseif O==2  % concentrations change vertically and repeats horizontally
    nn=1;
    nc=1;
    DD(1,2)=Cl(1);

  for ik=1:I-1
      if DD(ik+1,1)==DD(ik,1)
          DD(ik+1,2)=Cl(nc);
      else
          nn=nn+1;
          if nn>Nr
              nn=1;
              nc=nc+1;
              if nc>Conc
                  error('something wrong with concentrations arrangements')
              end
          end
          DD(ik+1,2)=Cl(nc);
      end
  end
end

BB=sortrows(DD,2);
save('data.txt', 'BB', '-ascii', '-tabs')
```

**Part 2. The Ratio method**

```
% Thermofluor data
% Part 2: Determining the melting points, Tm, and Gibbs energy using the "Ratio" method

% Inputs: cured by "MeltingOriginalData.m" main file melting data: "data.txt"
% Outputs: table of melting points, and graphs of all data


disp('--------------Starting the fitting------------------')

%-----------------------Parameter definitions-----------------------------------------
%Nr: number of repeats of each mutant
%Tbeg: lower limit of temperature for fitting
%Tend: upper limit of temperature for fitting
%scl: scaling parameter for the figures (e.g. dT=(Thigh-Tlow)/scl;T=[Tlow-dT;Tmax+dT])
%smrd: determines the degree of smoothing: rd=round(round(lm*smrd=0.1)/2)*2+1;G=smooth(G,rd);

%--------------------End of parameter definitions----------------------

clear all;
D=load('data.txt');
[dr,dc]=size(D);

ss=input('Would you like to discard some data (y/n)?','s');
Nwell=[];
if ss=='y' || ss=='Y'
    fid = fopen('Discards_R.txt', 'wt');
    while ss=='y' || ss=='Y'
        if ss~='y' && ss~='Y'
            break
        end
        dk = input('Well number=', 's');
        Nwell=[Nwell;str2double(dk)];
        Msg0=strcat('Well_#_', dk,'  was discarded manually at the beginning');%num2str(MM(jj)));
        fprintf(fid,'%s \n',Msg0);
        ss=input('More data (y/n)?','s');
    end
    fclose(fid);
end


Tbeg=27;Tend=95;% range of temperature for study

T0=29; %if Tmin will be found less then T0 degree then Tmin will be assigned by hand
T1=65; %if Tmin will be found more then T1 degree then Tmin will be assigned by hand

T2=70; %if Tmax will be found less then T2 degree then Tmax will be assigned by hand
T3=90; %if Tmax will be found more then T3 degree then Tmax will be assigned by hand


scl=20;
ki=0.1;
ke=0.1;
smrd=0.15;
Nr=6;% number of repeats of each mutant
thr=0.5;% share of points that should be on the right side of the melting curve


%------------------Saving Parameters-------------------------------------

fid = fopen('Parameters_R.txt','wt');
fprintf(fid,'%s \n \n','Fitting parameters:');
tl=['Nr ',' Tbeg ',' Tend ',' T0 ',' T1 ',' T2 ',' T3 ',' scl ',' ki ',' ke ',' smrd ',' thr'];
fprintf(fid,'%s \n',tl);
Tf=[Nr,Tbeg,Tend,T0,T1,T2,T3,scl,ki,ke,smrd,thr];
fprintf(fid,'%g \t %g \t %g \t %g \t %g \t %g \t %g \t %g \t %g \t %g \t %g \t %g \t \n \n \n',Tf');
fclose(fid);

%-------------------------------------------------


scrsz = get(0,'ScreenSize');%deterimning the screen size to pozition the figures

mkdir('Graphs_R');% creating directory for the output graphs
mkdir('Gibbs Energy_R');% creating directory for Gibbs energy files

Temp=D(:,3);% temperature

%limiting data to a sensable range of temperature
kT=find(Temp>Tbeg & Temp<Tend);


[MM idx] = unique( sort(D(:,1)) );
% MM is the set of well numbers
% idx is a vector of total number of data point up to current well

mm=[];% wells that will be included in the analyses
mt=[]; % mutants that will be included in the analyses
wellcounts = diff([0;idx]); % number of data points in each well

mw=length(MM); % number of wells for study
for ss=1:mw
    n=find(D(:,1)==MM(ss),1,'first');
    Mut(ss)=D(n,2);
end

Mut=Mut';
if size(Mut)~=size(MM)
    error('Something wrong with the data')
end


sc=0; % number of sets analized
nc=1;% index for the set for simulation
GminI=0; %indicates that Gmin is determined
GmaxI=0; %indicates that Gmax is determined

fid = fopen('Error logs_R.txt', 'wt'); % open file for error recording
errlog=0;% index for error: 0, no errors; 1, there are errors
Tmax01=[];
Tmin01=[];

%Giving a choice to manually input Tmin if not found or have it fixed
ss=input('Would you like to input common Tmin if not found by the program (y/n)?','s');
tm=[];
if ss=='y' || ss=='Y'
    tm = input('Tmin=', 's');
    Tmin=str2double(tm);

    if Tmin<T0 || Tmin>T1
                Tmin
                disp('You entered wrong Tmin. Please choose a correct value of Tmin manually')
                Tminm0 = input('Tmin=', 's');
                Tmin=str2double(Tminm0);
    end
end


for jj=1:mw % cycle for datasets
    disp('-----New dataset-----')
    abc=find(Nwell==MM(jj));

    if isempty(abc)
       xstr=int2str(MM(jj));% well number
       Vcon=int2str(Mut(jj));% mutant's number

       %reading the data from a given well (xx)

       Ig=find(D(:,1)==MM(jj));
       Ig=Ig(D(Ig,3)>Tbeg & D(Ig,3)<Tend);% limiting by temperature
       T=D(Ig,3);
       G=D(Ig,4);
       [m,n]=size(G);

       %minimum and maximum of the data for plotting purposes only so all
       %plots have the same scales
       Gmaximum=max(G);
       Gminimum=min(G);
       g=[0.9*Gminimum,1.1*Gmaximum];

       %----------Plotting------------------------------
       %---Plotting scales---
       Tlow=min(T);
       Thigh=max(T);
       dT=(Thigh-Tlow)/scl;
       Tlow=Tlow-dT;
       Thigh=Thigh+dT;
       Glow=min(G);
       Ghigh=max(G);
       dG=(Ghigh-Glow)/scl;
       Glow=Glow-dG;
       Ghigh=Ghigh+dG;
       %-------------

       %plotting original data
       figure('OuterPosition',[1 scrsz(4)/2 scrsz(3)/2 scrsz(4)/2])
       subplot(2,2,1)
       scatter(T,G,'.');
       title(['Well #:',xstr])
       axis([Tlow Thigh Glow Ghigh])
       hold on

      %------------------"Smoothing" the data for better presentation and finding peaks---------------------------------------------
      % remembering the raw data
      Gf=G;
      Tf=T;
      lm=length(G);

      % smoothing
      rd=round(round(lm*smrd)/2)*2+1;
      G=smooth(G,rd);

      %---Plotting the smoothed data----------
      subplot(2,2,3)
      scatter(T,G,'.');
      title('Smoothed data')
      axis([Tlow Thigh Glow Ghigh])
      hold on

      %----Finding local mins and maxs in the smoothed data-----
      [gmaximum,nmax0]=findpeaks(G);
      [gminimum,nmin0]=findpeaks(-G);

      a0=find(T(nmax0)>T2 & T(nmax0)<T3);
      nmax=nmax0(a0);
      a1=find(T(nmin0)>T0 & T(nmin0)<T1);
      nmin=nmin0(a1);

      if isempty(nmax) || length(nmax)>1

         disp('No reliable max has been found by "findpeaks". Please choose a value of Tmax manually (0 to skip)')
         Tmaxm0 = input('Tmax=', 's');
         Tmax01=str2double(Tmaxm0);

         if Tmax01==0
            errlog=1;
            errmessage=['Well #',xstr,'  was discarded because no max has been found by "findpeaks"']

         elseif Tmax01<T2 || Tmax01>T3
            Tmax01
            disp('You entered wrong Tmax. Please choose a correct value of Tmax manually (0 to skip)')
            Tmaxm0 = input('Tmax=', 's');
            Tmax01=str2double(Tmaxm0);
         end

         [xmax,nmax]=min(abs(T-Tmax01));
         gmaximum=G(nmax);
      end

      Tmax=T(nmax);

      if isempty(nmin) || length(nmin)>1
         if isempty(tm)
            disp('No reliable min has been found by "findpeaks". Please choose a value of Tmin manually (0 to skip)')
            Tminm0 = input('Tmin=', 's');
            Tmin01=str2double(Tminm0);

            if Tmin01==0
                errlog=1;
                errmessage=['Well #',xstr,'  was discarded because no min has been found by "findpeaks"']

            elseif Tmin01<T0 || Tmin01>T1
                Tmin01
                disp('You entered wrong Tmin. Please choose a correct value of Tmin manually')
                Tminm0 = input('Tmin=', 's');
                Tmin01=str2double(Tminm0);
            end
         else
             Tmin01=Tmin;
         end
        [xmin,nmin]=min(abs(T-Tmin01));
        gminimum=G(nmin);
      end
      Tmin=T(nmin);


      if errlog==0

            tmax=[T(nmax),T(nmax)];
            tmin=[T(nmin),T(nmin)];

            % nmax is the position (index) of the Tmax and Gmax
            % nmin is the position (index) of the Tmin and Gmin

            %---Plotting the data indicating the global max and min
            subplot(2,2,1)
            plot(tmin,g,'r')
            plot(tmax,g,'g')
            xlabel('')
            ylabel('Fluo Readings')
            axis([Tlow Thigh Glow Ghigh])
            hold on
            %--------------------------

            if nmax-nmin<5 %condition for not fitting due to insufficiant number of data points
               subplot(2,2,3)
               plot(tmin,g,'r')
               plot(tmax,g,'g')

               errlog=1;
               errmessage=['Well #',xstr,'  was discarded due to insufficiant points for fitting']
               disp('Err=2')
               disp('Push any button to continue')
               pause
            end

            [nT,kT]=size(T);
            [nG,kG]=size(G);

            %just for control
            if nG~=nT
               error('number of points on two axises are not equal')
            end


        %------------Finding dGibbs from line extrapolations and ratios-----
        if errlog==0
           sc=sc+1;
           % 1. Fitting the initial part by a straight line
           KiniI=ceil(nmin*ki);

        if KiniI==0
           KiniI=1;
        end

        KiniE=floor(nmin*(1-ki));
        if KiniE<5
           KiniE=5;
        end

        % estimating the slope
        Ti=T(KiniI:KiniE);
        Gi=G(KiniI:KiniE);

        dt=T(KiniI)-T(KiniE);
        dr=G(KiniI)-G(KiniE);
        b=dr/dt;
        a=G(KiniI)-b*T(KiniI);

        options = fitoptions('Method','NonlinearLeastSquares',...
               'Startpoint',[a b],...
               'MaxFunEvals',10000,...
               'MaxIter',10000);

        f = fittype('(a + b*x)','options',options);

        [c2,gof2] = fit(Ti,Gi,f);
        bb=c2.b;
        %---------------------

        [ai(sc),bi(sc)]=FitLines(Ti,Gi,1);

        if ai(sc)==0 && bi(sc)==0
           errlog=1;
           errmessage=['Well #',xstr,'  was discarded due to insufficiant points for fitting at the beginning']
           sc=sc-1;
        end

        klow=round((KiniE+KiniI)/2);
        Tlow=T(klow);

        % 2. Fitting the end part by a straight line

        KendI=nmax+round((lm-nmax)*ke);

        if KendI>lm
           KendI=lm-6;
        end

        KendE=lm-round((lm-nmax)*ke);
        if KendE<=nmax
           KendE=lm-1;
        end

        % estimating the slope
        Te=T(KendI:KendE);
        Ge=G(KendI:KendE);

        dt=T(KendI)-T(KendE);
        dr=G(KendI)-G(KendE);
        b=dr/dt;
        a=G(KendI)-b*T(KendI);

        options = fitoptions('Method','NonlinearLeastSquares',...
               'Startpoint',[a b],...
               'MaxFunEvals',10000,...
               'MaxIter',10000);

        f = fittype('(a + b*x)','options',options);

        [c2,gof2] = fit(Te,Ge,f);

        ae(sc)=c2.a;
        be(sc)=c2.b;

        khigh=round((KendE+KendI)/2);
        Thigh=T(khigh);

        %-----Simulating the ini and end part by lines and plotting them
        if errlog==0
           TdG=T(klow:khigh);
           GdG=G(klow:khigh);
           Gini=ai(sc)+bi(sc)*TdG;
           Gend=ae(sc)+be(sc)*TdG;

           subplot(2,2,1)
           plot(TdG,Gini,'r','LineWidth',2)
           plot(TdG,Gend,'g','LineWidth',2)
           hold on

           subplot(2,2,3)
           plot(TdG,Gini,'r','LineWidth',2)
           plot(TdG,Gend,'g','LineWidth',2)
           hold on
           %------------------------------------------------

           TdG=[];
           GdG=[];
           Keq=[];
           Gibbs=[];
           TdG=T(nmin:nmax);
           GdG=Gf(nmin:nmax);
           Gini=ai(sc)+bi(sc)*TdG;
           Gend=ae(sc)+be(sc)*TdG;
           Kend=Gend-GdG;
           Kini=GdG-Gini;
           Nend=length(Kend);
           Nini=length(Kini);
           AP=find(Kend>0);
           AN=find(Kini>0);
           NP=length(AP);
           NN=length(AN);


           if NP/Nend<thr || NN/Nini<thr || isempty(NP) || isempty(NN)
              NbyNend=NP/Nend
              NbyNini=NN/Nini
              errlog=1;
              errmessage=['Well #',xstr,'  was discarded due to bad data at the beginning or end']
              sc=sc-1;
              disp('Push any button to continue')
              pause
           end

           if errlog==0
              Keq=(Gend-GdG)./(GdG-Gini);
              Gibbs=1.99*(TdG+273).*log(Keq);

              %---------Finding Tm------------------------------
              %The condition for Tm is Keq=1 or ln(Keq)=0 and Gibbs=0
              [GP,x]=find(Gibbs>0,1,'last');
              [GN,y]=find(Gibbs<0,1,'first');

              bem=(Gibbs(GP)-Gibbs(GN))/(TdG(GP)-TdG(GN));
              aem=Gibbs(GN)-bem*TdG(GN);
              Tm(sc)=-aem/bem;
              mm(sc)=MM(jj);
              mut(sc)=Mut(jj);

           end


           dG(sc)=G(T==tmax(1,1))-G(T==tmin(1,1)); %"amplitude" of the melting
           Amp(sc)=dG(sc);

           if dG(sc)<100
              disp('Small amplitude')
              Ampl=dG(sc)
              errlog=1; % data set will be discarded because of small amplitude of the signal
              pause
           end

           %-----------Final Plotting------------------
            subplot(2,2,1)
            plot([Tm(sc),Tm(sc)],g,'k','LineWidth',2)
            xlabel('Temperature')
            ylabel('Fluo Readings')
            hold on

            subplot(2,2,2)
            title('Gibbs energy')
            scatter(TdG,Gibbs,'.b');
            title(['Mutant=',Vcon])
            hold on
            plot([Tm(sc),Tm(sc)],[min(Gibbs),max(Gibbs)],'k','LineWidth',2)
            xlabel('Temperature')
            ylabel('Gibbs energy')
            axis auto
            hold on

            subplot(2,2,3)
            plot(tmin,g,'r')
            hold on
            plot(tmax,g,'g')
            hold on
            plot([Tm(sc),Tm(sc)],g,'k','LineWidth',2')
            xlabel('Temperature')
            ylabel('Fluo Readings')
            axis([Tlow Thigh Glow Ghigh])
            hold on

            subplot(2,2,4)
            scatter(1./(TdG+273),Gibbs,'.b');
            hold on
            plot([1/(Tm(sc)+273),1/(Tm(sc)+273)],[min(Gibbs),max(Gibbs)],'k','LineWidth',2)
            xlabel('1/Temperature')
            ylabel('Gibbs energy')
            hold off
        end
        end
      end


      if errlog~=0
         disp('saving error messages')
         fprintf(fid,'%s \n',errmessage);
         disp('Push any button to continue')
         pause
      elseif errlog==0
         ffile=strcat('Gibbs Energy_R/Gibbs_R_Well_', num2str(MM(jj)),'_Conc_',Vcon,'.txt');
         GibbsData=[TdG,Gibbs];
         save(ffile, 'GibbsData','-ascii', '-tabs'); % saving the Gibbs energy for a given well
      end

      ffig=strcat('Well #', num2str(MM(jj)),'_Conc_',Vcon);
      saveas(gcf,['Graphs_R',filesep,ffig],'jpg') % saving the figure of simulation
    end

    errlog=0;
    GminI=0;
    GmaxI=0;

end

fclose(fid);

%-----------------End of simulation------------------------


%-------------------Saving results------------
Fnm1=strcat('Ratio_results.xls');

R={'Well #','Mutant Name','Mutant','Tm'};
Tmelting=[mm',mut',mut',Tm'];

ss=input('Would you like to discard some data (y/n)?','s');
Nwell=[];
if ss=='y' || ss=='Y'
    fid = fopen('Discards_R.txt', 'wt');
    while ss=='y' || ss=='Y'
        if ss~='y' && ss~='Y'
            break
        end
        dk = input('Well number=', 's');
        Nwell=str2double(dk);
        melt=find(Tmelting(:,1)==Nwell);
        Tmelting(melt,:)=[];
        Msg0=strcat('Well_#_', dk,'  was discarded manually');%num2str(MM(jj)));
        fprintf(fid,'%s \n',Msg0);
        ss=input('More data (y/n)?','s');

    end
    fclose(fid);
end


T=num2cell(Tmelting);

for ii=1:length(T)
    Mutant=mutants(Mut(ii));
    MutN(ii)={Mutant};
end

MutN=MutN';
Tt=T;
Tt(:,2)=MutN;


d=[R;Tt];
s=xlswrite(Fnm1,d,1)

R={'Well #','Mutant Name','Mutant','Tm'};

ds=dataset('xlsfile',Fnm1);
Mutnt=grpstats(ds.Mutant,ds.Mutant);

mTm=grpstats(ds.Tm,ds.Mutant);
sdevTm=grpstats(ds.Tm,ds.Mutant,'std');
serrTm=grpstats(ds.Tm,ds.Mutant,'sem');

nmbr=grpstats(ds.Tm,ds.Mutant,'numel');

Rtt={'Mutant',' Tm ','stddevTm','serrTm','number of repeats'};


k=length(Mutnt);
Rxx=[];
Rss=[];

for i=1:k
    mutt=mutants(Mutnt(i));
    AMutant(i)={mutt};
    Rxx={mTm(i),sdevTm(i),serrTm(i),nmbr(i)};
    Rss=[Rss;Rxx];
end

Rss=[AMutant',Rss];

Rmean=[Rtt;Rss];
s=xlswrite(Fnm1,Rmean,2)
```

**Part 3. Thermodynamic model**

```
% Thermofluor data
% Part 3: Determining the melting points, Tm, and Gibbs energy using the "Model" method

% Inputs: cured by "MeltingOriginalData.m" main file melting data: "data.txt"
% Outputs: table of melting points, fitting scores, and graphs of all data
% with the fitted curves

disp('--------------Starting the fitting------------------')

%-----------------------Parameter definitions-----------------------------------------
%Nr: number of repeats of each Mutant
%Mutant: set of mutants that should be fitted
%Tbeg: lower limit of temperature for fitting
%Tend: upper limit of temperature for fitting
%scl: scaling parameter for the figures (e.g. dT=(Thigh-Tlow)/scl;T=[Tlow-dT;Tmax+dT])
%kiI: determines the first point of the beginning straight line
%kiE: determines the last point of the beginning straight line
%keI: determines the first point of the ending straight line
%keE: determines the last point of the ending straight line
%smrd: determines the degree of smoothing: rd=round(round(lm*smrd=0.1)/2)*2+1;G=smooth(G,rd);
%mthd: choose a fitting method from the list: 'Levenberg-Marquardt','Gauss-Newton', or 'Trust-Region'. The default is 'Trust-Region'

%--------------------End of parameter definitions----------------------

clear all;
D=load('data.txt');
[dr,dc]=size(D);

% Some data might be not good.  Here is the place to get rid of it
ss=input('Would you like to discard some data (y/n)?','s');
Nwell=[];
if ss=='y' || ss=='Y'
    fid = fopen('Discards_M.txt', 'wt');
    while ss=='y' || ss=='Y'
        if ss~='y' && ss~='Y'
            break
        end
        dk = input('Well number=', 's');
        Nwell=[Nwell;str2double(dk)];
        Msg0=strcat('Well_#_', dk,'  was discarded manually at the beginning');%num2str(MM(jj)));
        fprintf(fid,'%s \n',Msg0);
        ss=input('More data (y/n)?','s');
    end
    fclose(fid);
end


R=1.99;% Gas constant: R=1.99 cal/mol/K

Tbeg=27;Tend=95;% range of temperature for study

T0=30; %if Tmin will be found less then T0 degree then Tmin will be assigned by hand
T1=65; %if Tmin will be found more then T1 degree then Tmin will be assigned by hand

T2=70; %if Tmax will be found less then T2 degree then Tmax will be assigned by hand
T3=90; %if Tmax will be found more then T3 degree then Tmax will be assigned by hand


scl=20;
ki=0.1;
ke=0.1;
smrd=0.15;
mthd='Trust-Region';
Nr=6;% number of repeats of each mutant


%------------------Saving Parameters-------------------------------------

fid = fopen('Parameters_M.txt','wt');
fprintf(fid,'%s \n \n','Fitting parameters:');
fprintf(fid,'%s \n','Fitting method:');
fprintf(fid,'%s \n \n',mthd);
tl=['Nr ',' Tbeg ',' Tend ',' T0 ',' T1 ',' T2 ',' T3 ',' scl ',' ki ',' ke ',' smrd '];
fprintf(fid,'%s \n',tl);
Tf=[Nr,Tbeg,Tend,T0,T1,T2,T3,scl,ki,ke,smrd];
fprintf(fid,'%g \t %g \t %g \t %g \t %g \t %g \t %g \t %g \t %g \t %g \t %g \t \n \n \n',Tf');
fclose(fid);

%-------------------------------------------------


scrsz = get(0,'ScreenSize');%deterimning the screen size to pozition the figures

mkdir('Graphs_M');% creating folder for the graphs
mkdir('Gibbs Energy_M');% creating folder for Gibbs energies

Temp=D(:,3);% temperature

%limiting data to a reasonable range of temperature
kT=find(Temp>Tbeg & Temp<Tend);


[MM idx] = unique( sort(D(:,1)) );
% MM is the set of well numbers
% idx is a vector of total number of data point up to current well

mm=[];% wells that will be included in the analyses
mt=[]; % wells that will be included in the analyses
wellcounts = diff([0;idx]); % number of data points in each well

mw=length(MM); % number of wells for study

%creating list of mutant numbers
for ss=1:mw
    n=find(D(:,1)==MM(ss),1,'first');
    Mut(ss)=D(n,2);
end

% control
Mut=Mut';
if size(Mut)~=size(MM)
    error('Something wrong with the data')
end


%---------------------

sc=0; % number of sets analized

fid = fopen('Error logs_M.txt', 'wt');
errlog=0;
Tmax01=[];
Tmin01=[];

%Giving a choice to manually input Tmin if not found or have it fixed
ss=input('Would you like to input common Tmin if not found by the program (y/n)?','s');
tm=[];
if ss=='y' || ss=='Y'
    tm = input('Tmin=', 's');
    Tmin=str2double(tm);

    if Tmin<T0 || Tmin>T1
                Tmin
                disp('You entered wrong Tmin. Please choose a correct value of Tmin manually')
                Tminm0 = input('Tmin=', 's');
                Tmin=str2double(Tminm0);
    end
end


%----------------Starting the simulation------------------------
for jj=1:mw % dataset
        disp('-----New dataset-----')
        abc=find(Nwell==MM(jj));

        if isempty(abc)
           xstr=int2str(MM(jj));% well number
           Vcon=int2str(Mut(jj));% mutant's number

           %reading the data from a given well (xx)

            Ig=find(D(:,1)==MM(jj));

            Ig=Ig(D(Ig,3)>Tbeg & D(Ig,3)<Tend);% limiting by temperature
            T=D(Ig,3);

            G=D(Ig,4);
            [m,n]=size(G);

            %minimum and maximum of the data for plotting purposes only so all
            %plots have the same scales
            Gmaximum=max(G);
            Gminimum=min(G);
            g=[0.9*Gminimum,1.1*Gmaximum];

            %---Plotting scales---
            Tlow=min(T);
            Thigh=max(T);
            dT=(Thigh-Tlow)/scl;
            Tlow=Tlow-dT;
            Thigh=Thigh+dT;
            Glow=min(G);
            Ghigh=max(G);
            dG=(Ghigh-Glow)/scl;
            Glow=Glow-dG;
            Ghigh=Ghigh+dG;
            %-------------

            %plotting original data
            figure('OuterPosition',[1 scrsz(4)/2 scrsz(3)/2 scrsz(4)/2])
            subplot(2,2,1)
            scatter(T,G,'.');
            title(['Well #:',xstr])
            axis([Tlow Thigh Glow Ghigh])
            hold on

            %------------------"Smoothing" the data for better presentation and finding peaks---------------------------------------------
            % remembering the raw data
            Gf=G;
            Tf=T;
            lm=length(G);
            % smoothing
            rd=round(round(lm*smrd)/2)*2+1;
            G=smooth(G,rd);

            %---Plotting the smoothed data----------
            subplot(2,2,3)
            scatter(T,G,'.');
            title('Smoothed data')
            axis([Tlow Thigh Glow Ghigh])
            hold on

            %----Finding local mins and maxs in the smoothed data-----
            [gmaximum,nmax0]=findpeaks(G);
            [gminimum,nmin0]=findpeaks(-G);

            a0=find(T(nmax0)>T2 & T(nmax0)<T3);
            nmax=nmax0(a0);
            a1=find(T(nmin0)>T0 & T(nmin0)<T1);
            nmin=nmin0(a1);

            % checking whether maxima are found and if not input manually
            if isempty(nmax) || length(nmax)>1

               disp('No reliable max has been found by "findpeaks". Please choose a value of Tmax manually (0 to skip)')
               Tmaxm0 = input('Tmax=', 's');
               Tmax01=str2double(Tmaxm0);

               if Tmax01==0
                   errlog=1;
                   errmessage=['Well #',xstr,'  was discarded because no max has been found by "findpeaks"']

               elseif Tmax01<T2 || Tmax01>T3
                   Tmax01
                   disp('You entered wrong Tmax. Please choose a correct value of Tmax manually (0 to skip)')
                   Tmaxm0 = input('Tmax=', 's');
                   Tmax01=str2double(Tmaxm0);
               end

               [xmax,nmax]=min(abs(T-Tmax01));
               gmaximum=G(nmax);
            end

            Tmax=T(nmax);

             % checking whether minima are found and if not input manually
             if isempty(nmin) || length(nmin)>1
                if isempty(tm)
                   disp('No reliable min has been found by "findpeaks". Please choose a value of Tmin manually (0 to skip)')
                   Tminm0 = input('Tmin=', 's');
                   Tmin01=str2double(Tminm0);

                   if Tmin01==0
                       errlog=1;
                       errmessage=['Well #',xstr,'  was discarded because no min has been found by "findpeaks"']

                   elseif Tmin01<T0 || Tmin01>T1
                       Tmin01
                       disp('You entered wrong Tmin. Please choose a correct value of Tmin manually')
                       Tminm0 = input('Tmin=', 's');
                       Tmin01=str2double(Tminm0);
                   end
                else
                Tmin01=Tmin;
                end

                [xmin,nmin]=min(abs(T-Tmin01));
                gminimum=G(nmin);
             end

             Tmin=T(nmin);


             if errlog==0
                tmax=[T(nmax),T(nmax)];
                tmin=[T(nmin),T(nmin)];

                % nmax is the position (index) of the Tmax and Gmax
                % nmin is the position (index) of the Tmin and Gmin

                %---Plotting the data indicating the global max and min
                subplot(2,2,1)
                plot(tmin,g,'r')
                plot(tmax,g,'g')
                xlabel('')
                ylabel('Fluo Readings')
                axis([Tlow Thigh Glow Ghigh])
                hold on
                %--------------------------


                if nmax-nmin<5 %condition for not fitting due to insufficiant number of data points
                    subplot(2,2,3)
                    plot(tmin,g,'r')
                    plot(tmax,g,'g')

                    errlog=1;
                    errmessage=['Well #',xstr,'  was discarded due to insufficiant points for fitting']
                    disp('Push any button to continue')
                    pause
                end

                [nT,kT]=size(T);
                [nG,kG]=size(G);

                %just for control
                if nG~=nT
                   error('number of points on two axises are not equal')
                end

             end


            %-------Fitting data by formula-----------------------
            if errlog==0
               sc=sc+1;

               % 1. Estimating the slope and intercept of the initial part by straight line
                KiniI=ceil(nmin*ki);

                if KiniI==0
                   KiniI=1;
                end

                KiniE=floor(nmin*(1-ki));

                if KiniE<5
                   KiniE=5;
                end


                Ti=T(KiniI:KiniE);
                Gi=G(KiniI:KiniE);

                dt=T(KiniI)-T(KiniE);
                dr=G(KiniI)-G(KiniE);
                b=dr/dt;
                a=G(KiniI)-b*T(KiniI);
                ai(sc)=a;
                bi(sc)=b;


                if ai(sc)==0 || bi(sc)==0 || isempty(ai(sc)) || isempty(bi(sc))
                   errlog=1;
                   errmessage=['Well #',xstr,'  was discarded due to insufficiant points for fitting at the beginning']
                   sc=sc-1;
                end

                klow=round((KiniE+KiniI)/2);
                Tlow=T(klow);
                %--------------------------------------------------------


                % 2. Estimating the slope and intercept of the end part by straight line

                if errlog==0

                   KendI=nmax+round((lm-nmax)*ke);

                   if KendI>lm
                      KendI=lm-6;
                   end

                   KendE=lm-round((lm-nmax)*ke);
                   if KendE<=nmax
                      KendE=lm-1;
                   end


                   Te=T(KendI:KendE);
                   Ge=G(KendI:KendE);

                   dt=T(KendI)-T(KendE);
                   dr=G(KendI)-G(KendE);
                   b=dr/dt;
                   a=G(KendI)-b*T(KendI);

                   ae(sc)=a;
                   be(sc)=b;

                   if ae(sc)==0 || be(sc)==0 || isempty(ae(sc)) || isempty(be(sc))
                      errlog=1;
                      errmessage=['Well #',xstr,'  was discarded due to insufficiant points for fitting at the end']
                      sc=sc-1;
                   end

                   khigh=round((KendE+KendI)/2);
                   Thigh=T(khigh);

                   %--------------------------------------------------------


                    %Actual fitting
                    if errlog==0

                       TdG=T(klow:khigh);
                       GdG=G(klow:khigh);

                       G=Gf;% back from smoothed data to the original

                       To=(Tmin+Tmax)/2;

                       dCp(sc)=3000;% for the fitting we use dCp fixed at 3000 cal/Celcius

                       options = fitoptions('Method','NonlinearLeastSquares',...
                                'Startpoint',[ai(sc) bi(sc) ae(sc) be(sc) 70000 To],...
                                'MaxFunEvals',100000,...
                                'MaxIter',100000,...
                                'Algorithm',mthd);

                        f = fittype('(a+b*x)+ (c+d*x)/(1.0 + exp((e*(1-(x+273)/(g+273))-3000*((g-x)+(x+273)*log((x+273)/(g+273))))/(x+273)/1.99))','options',options);

                        [c2,gof2] = fit(Tf,Gf,f);

                        %Fitting parameters
                        y0i=c2.a;
                        bi=c2.b;
                        y0e=c2.c;
                        be=c2.d;
                        dH(sc)=c2.e;
                        Tm(sc)=c2.g;

                        subplot(2,2,1)
                        plot(c2,'m')
                        legend off
                        hold on

                        subplot(2,2,3)
                        plot(c2,'m')
                        legend off
                        hold on
                    end

                    if Tm(sc)<Tmin || Tm(sc)>Tmax
                       errlog=1;
                       errmessage=['Set #',xstr,'  was discarded due to Tm being beyond limits']
                       disp('push any button to continue')
                       pause
                    end

                    if errlog==0

                        %Fitting statistics
                        sse=gof2.sse;% Sum squared error performance function
                        rsquare=gof2.rsquare;% Coefficient of determination
                        dfe=gof2.dfe;% Degrees of freedom
                        adjrsquare=gof2.adjrsquare;%Degree-of-freedom adjusted coefficient of determination
                        rmse=gof2.rmse;% Root mean squared error (standard error)

                        mm(sc)=MM(jj);
                        mt(sc)=Mut(jj);

                        Gibbs=(dH(sc)*(1-(TdG+273)/(Tm(sc)+273))-dCp(sc)*((Tm(sc)-TdG)+(TdG+273).*log((TdG+273)/(Tm(sc)+273))));
                        P1=1./(1+exp(-Gibbs./R./(TdG+273)));% probability of state "1"
                        P2=1./(1+exp(Gibbs./R./(TdG+273)));% probability of state "2"


                        %-----------Final Plotting------------------
                        subplot(2,2,1)
                        plot([Tm(sc),Tm(sc)],g,'k','LineWidth',2)
                        xlabel('Temperature')
                        ylabel('Fluo Readings')
                        hold on

                        subplot(2,2,2)
                        title('Gibbs energy (a.u.)')
                        scatter(TdG,Gibbs,'.b');
                        title(['Mutant=',Vcon])
                        hold on

                        plot([Tm(sc),Tm(sc)],[min(Gibbs),max(Gibbs)],'k','LineWidth',2)
                        xlabel('Temperature')
                        ylabel('Gibbs energy, cal')
                        axis auto
                        hold on

                        subplot(2,2,3)
                        plot(tmin,g,'r')
                        hold on
                        plot(tmax,g,'g')
                        hold on

                        plot([Tm(sc),Tm(sc)],g,'k','LineWidth',2')
                        xlabel('Temperature')
                        ylabel('Fluo Readings')
                        axis([Tlow Thigh Glow Ghigh])
                        hold on

                        subplot(2,2,4)
                        scatter(TdG,P1,'.b');
                        hold on
                        scatter(TdG,P2,'.r');
                        xlabel('Temperature')
                        ylabel('Probabilities')
                        hold off
                    end
                end
            end

            if errlog~=0
                disp('saving error messages')
                errlog=errlog
                fprintf(fid,'%s \n',errmessage);
                disp('Push any button to continue')
                pause
            elseif errlog==0
                ffile=strcat('Gibbs Energy_M/Gibbs_M_Well_', num2str(MM(jj)),'_Conc_',Vcon,'.txt');
                GibbsData=[TdG,Gibbs];
                save(ffile, 'GibbsData','-ascii', '-tabs'); % saving the Gibbs energy for a given well
            end

            ffig=strcat('Well #', num2str(MM(jj)),'_Conc_',Vcon);
            saveas(gcf,['Graphs_M',filesep,ffig],'jpg')
        end

        errlog=0;

end

fclose(fid);

%-----------------End of analysis------------------------


%----------Saving results------------
Fnm1=strcat('Model_results.xls');

R={'Well #','Mutant Name','Mutant','Tm'};
Tmelting=[mm',mt',mt',Tm'];

ss=input('Would you like to discard some data (y/n)?','s');% A chance to discard bad data and/or fitting
Nwell=[];

if ss=='y' || ss=='Y'
    fid = fopen('Discards_R.txt', 'wt');

    while ss=='y' || ss=='Y'
        if ss~='y' && ss~='Y'
            break
        end
        dk = input('Well number=', 's');
        Nwell=str2double(dk);
        melt=find(Tmelting(:,1)==Nwell);
        Tmelting(melt,:)=[];
        Msg0=strcat('Well_#_', dk,'  was discarded manually');%num2str(MM(jj)));
        fprintf(fid,'%s \n',Msg0);
        ss=input('More data (y/n)?','s');

    end
    fclose(fid);
end


T=num2cell(Tmelting);

for ii=1:length(T)
    Mutant=mutants(Mut(ii));
    MutN(ii)={Mutant};
end

MutN=MutN';
Tt=T;
Tt(:,2)=MutN;


d=[R;Tt];
s=xlswrite(Fnm1,d,1) %saving results of fitting (all sets)

R={'Well #','Mutant Name','Mutant','Tm'};

ds=dataset('xlsfile',Fnm1);
Mutnt=grpstats(ds.Mutant,ds.Mutant);

mTm=grpstats(ds.Tm,ds.Mutant);
sdevTm=grpstats(ds.Tm,ds.Mutant,'std');
serrTm=grpstats(ds.Tm,ds.Mutant,'sem');

nmbr=grpstats(ds.Tm,ds.Mutant,'numel');

Rtt={'Mutant',' Tm ','stddevTm','serrTm','number of repeats'};


k=length(Mutnt);
Rxx=[];
Rss=[];

for i=1:k
    mutt=mutants(Mutnt(i));
    AMutant(i)={mutt};
    Rxx={mTm(i),sdevTm(i),serrTm(i),nmbr(i)};
    Rss=[Rss;Rxx];
end

Rss=[AMutant',Rss];

Rmean=[Rtt;Rss];
s=xlswrite(Fnm1,Rmean,2) %saving results of fitting (only means)
```

**Functions**

*Fitlines*

```
%function [a,b]=FitLines(Tb,Rb,Sg,ik)
function [a,b]=FitLines(Tb,Rb,ik)
% Function to fit one of the three parts of ThermaFluor data by three straight lines
% The requirements for the fittings are:
% 1. Include maximum number of data points from a given interval
% 2. Find the parameters that include top 25% best fits (using "rsquare" as
% a criteria)

% Sg gives the sign of the slope
% ik=1: initial part
% ik=3: end part

Tbm=mean(Tb); % mean value of temperature range
[dTb,kTb]=min(abs(Tb-Tbm)); % the distance between and position of the mean and closest data point
%kTb=find(round((Tbm-dTb)*10)==round(Tb*10) | round((Tbm+dTb)*10)==round(Tb*10), 1 ); %finding the position of the lowest closest to the mean point
[rb,cb]=size(Tb);

if rb<5 % if there is not enough points for fitting go back to program and assign a=0 and b=0
    a=0;
    b=0;
    return
end

km=min((rb-kTb),kTb); % finding the distance from the mean to the closest end
%km=min(floor(rb-kTb),kTb)-1; % finding the distance from the mean to the closest end

ap=0;% number of positive slopes
an=0;% number of negative slopes
a0=0;% number of zero slopes

for ii=1:km-1
Tf=[];
Gf=[];
k1=kTb-ii;% moving "down" from mean
k2=kTb+ii;% moving "up" from mean
Tf=Tb(k1:k2);
Rf=Rb(k1:k2);
%{
if abs(k2-k1)<2
    Nf(ii,1)=ii-1;
    Nf(ii,2)=Nf(ii-1,2);
    y0b(ii)=y0b(ii-1);
    bb(ii)=bb(ii-1);
    rsquare(ii)=rsquare(ii-1);
    break
end
%}
dt=Tb(k1)-Tb(k2);
dr=Rb(k1)-Rb(k2);
b=dr/dt; % estimating the slope
a=Rb(k1)-b*Tb(k1); % estimating the intersection
options = fitoptions('Method','NonlinearLeastSquares',...
               'Startpoint',[a b],...
               'MaxFunEvals',10000,...
               'MaxIter',10000);

f = fittype('(a + b*x)','options',options);

[c2,gof2] = fit(Tf,Rf,f);

y0b(ii)=c2.a;
bb(ii)=c2.b;

if bb(ii)>0
    ap=ap+1;
elseif bb(ii)<0
    an=an+1;
else
    a0=a0+1;
end


%Fitting statistics
sse(ii)=gof2.sse;% Sum squared error performance function
rsquare(ii)=gof2.rsquare;% Coefficient of determination
dfe(ii)=gof2.dfe;% Degrees of freedom
adjrsquare(ii)=gof2.adjrsquare;%Degree-of-freedom adjusted coefficient of determination
rmse(ii)=gof2.rmse;% Root mean squared error (standard error)
Nf(ii,1)=ii;
Nf(ii,2)=length(Tf);
end

at=ap+an+a0; %total number of slopes

if at~=ii
    disp('There is an error in the fitting of the linear part')
    a=0;
    b=0;
    pause
    disp('Push any button')
    return
end


if ap>2/3*at
   Sg=1;
elseif an>2/3*at
    Sg=-1;
elseif a0>2/3*at
    Sg=0;
else
    disp('It is impossible to determine the sign of the slope')
    a=0;
    b=0;
    pause
    disp('Push any button')
    return
end


Mtr=[Nf,rsquare',y0b',bb'];
[rw,cl]=size(Mtr);
if Sg>0 || Sg<0
    [al,bn]=find(Sg*Mtr(:,5)>0);% slopes of right signs
elseif Sg==0
    [al,bn]=find(Mtr(:,5)==0);% no slopes
end

%{
if ik==1 % only for initial parts
    if isempty(al) || length(al)<0.3*rw
        Sg=-Sg;
        [al,bn]=find(-Sg*Mtr(:,5)<=0);% slopes of right signs
    end
end
%}

Mtr=Mtr(al,:);
bb=Mtr(:,5);

bbMed=median(bb);% median value of the slopes
bbMin=min(bb); % minimum of slope values
bbMax=max(bb); % maximum of slope values
db=min(abs(bbMed-bbMin),abs(bbMed-bbMax)); % choosing the smaller value around the median

xbb=[bbMed-db,bbMed+db];
ybb=[bbMed-db,bbMed+db];

Mtr1=Mtr(bb>bbMed-db & bb<bbMed+db,:);

erSq=Mtr1(:,3);
erSq75=max(erSq)*0.75;
n=find(erSq>=erSq75);
[inx,lx]=max(Mtr1(n,2));
a=Mtr1(n(lx),4);
b=Mtr1(n(lx),5);

if length(a)>1 || length(b)>1
    disp('The fitting of the linear part did not reveal a unique solution')
    a=0;
    b=0;
    pause
    disp('Push any button')
elseif isempty(a) || isempty(b)
    disp('The fitting of the linear part did not reveal any solutions')
    a=0;
    b=0;
    pause
    disp('Push any button')
end
return
```

*Mutants*

```
function Mutant=mutants(Mut)
%----------------Definition of mutant and ligand types------------------------
if Mut==0
    Mutant='WT';
elseif Mut==1
    Mutant='I4V';
elseif Mut==2
    Mutant='F26L';
elseif Mut==3
    Mutant='Y33F';
elseif Mut==4
    Mutant='F37I';
elseif Mut==5
    Mutant='I4V_F26L';
elseif Mut==6
    Mutant='I4V_Y33F';
elseif Mut==7
    Mutant='I4V_F37I';
elseif Mut==8
    Mutant='F26L_Y33F';
elseif Mut==9
    Mutant='F26L_F37I';
elseif Mut==10
    Mutant='Y33F_F37I';
elseif Mut==11
    Mutant='I4V-F26L-Y33F';
elseif Mut==12
    Mutant='I4V-F26L-F37I';
elseif Mut==13
    Mutant='I4V-Y33F-F37I';
elseif Mut==14
    Mutant='F26L-Y33F-F37I';
elseif Mut==15
    Mutant='I4V-F26L-Y33F-F37I';
end

return
```
